# Effects of different immunomodulating liposome-based adjuvants and injection sites on immunogenicity in pigs

**DOI:** 10.1101/2023.06.08.544239

**Authors:** Evelína Šťastná, Gitte Erbs, Kerstin Skovgaard, Jeanne Toft Jakobsen, Gabriel Kristian Pedersen, Gregers Jungersen

**Affiliations:** Infectious Disease Immunology, Centre for Vaccine Research, Statens Serum Institut, Copenhagen S, Denmark; Department of Biotechnology and Biomedicine, Technical University of Denmark, Kongens Lyngby, Denmark

**Keywords:** Pig, Immunization, Adjuvant, Injection site, Immunomodulation, Adaptive immunity

## Abstract

Vaccine adjuvants are able to boost immune responses and steer immunity towards a desired direction. Liposome-based cationic adjuvant formulations (CAFs) are effective in inducing cell-mediated immune responses in mice, non-human primates and humans. In the translation from mouse to humans, pigs could play an important role. In this study, we thus used commercial pigs housed under field conditions to investigate the effects of four different CAFs incorporating distinct immunomodulators: C-type lectin receptor ligands trehalose-6,6’-dibehenate and monomycolyl glycerol, toll-like receptor ligand Poly(I:C) or retinoic acid. The vaccines were formulated with a recombinant *Chlamydia* model protein antigen and administered via distinct injection routes. All adjuvants significantly increased antigen-specific IgA and IgG in serum, compared to non-adjuvanted antigen. Administering the vaccines through intramuscular and intraperitoneal routes induced significantly higher antigen-specific IgA and IgG serum antibodies, than the perirectal route. Although the immunizations triggered cell-mediated immunity, no significant differences between the adjuvants or injection sites were detected by intracellular flow cytometry or cytokine-release assays. Genes depicting T cell subtypes were monitored by qPCR, which revealed minor differences only. Our findings suggest that the adjuvant-specific signature of the tested adjuvant immunomodulation does not translate well from mice to pigs. This study provides new insights into immune responses to CAFs in the pig model, and highlights that adjuvant studies should be ideally carried out in the intended species of interest.

## 1. Introduction

Adjuvants are defined as vaccine components able to enhance and shape immune responses. Addition of an appropriate adjuvant fills several gaps in modern vaccine development, for instance by lowering the antigen (Ag) dose, reducing number of immunizations, or steering the immune responses e.g. by stimulating specific subsets of T cells [1], [2]. Choosing a suitable adjuvant is therefore an important step in vaccine design and development. With the current trend in rational vaccine design using recombinant, highly purified Ags with a tremendous safety level, but relatively low immunogenicity, compared to live attenuated or crude inactivated vaccine preparations, there is additional need for adjuvants to stimulate protective immunity [2], [3]. Currently, most adjuvants induce particularly humoral immunity, but for many diseases, versatile immune responses, including enhanced CD4^+^ and CD8^+^ T cell responses are required for protection [3], [4]. Cationic liposomes, comprised of phospholipid bilayers, represent a class of adjuvants also serving as a delivery system, because they can directly incorporate Ags or immunomodulatory molecules [2]. Cationic adjuvant formulations (CAFs) based on dimethyldioctadecylammonium (DDA) formulated into liposomes, are versatile in the ability to incorporate different immunomodulators, allowing the vaccine-induced immune response to be skewed towards a preferred response [5]. In contrast to emulsion adjuvants which drain quickly to the lymphatics, CAF liposomes remain at the injection site and rely on Ag-presenting cell (APC) uptake and migration of the activated APCs to the draining lymph node [6].

The first-developed adjuvant in the CAF family was CAF01, consisting of amphiphilic DDA surfactant, and stabilized by trehalose-6,6’-dibehenate (TDB) glycolipid with immunostimulatory effects [7]. The key receptor for TDB is the C-type lectin receptor Mincle [8]. In mice, TDB was shown to selectively activate innate immunity through APC signaling pathway FcRγ – Syk – Card9, directing towards the induction of a mixed T helper 1 and 17 (Th1/Th17) response [9]. CAF01 has previously been used in a prime-boost strategy for vaccination against *Chlamydia* in several settings, including Göttingen minipigs [10], non-human primates [11] and recently in a human clinical trial [12]. Moreover, CAF01 was tested as a BCG booster immunization in non-human primate tuberculosis model [13]. CAF09b, CAF16 and CAF23 are novel members of the CAF adjuvant platform, all based on the DDA liposome backbone [5]. CAF09b is unique in inducing cytotoxic CD8^+^ T cell responses following intraperitoneal (i.p.) injection [14]. CAF09b liposomes incorporate stabilizing glycolipid monomycolyl glycerol (MMG), instead of TDB, and immunomodulator Poly(I:C), acting like a TLR3 agonist [14]. CAF09b adjuvant is currently undergoing phase II clinical trial (NCT05309421), for treatment of metastatic melanoma, where it serves as an adjuvant inducing CD8^+^ T cell responses in a personalized neoantigen peptide-based vaccine [15], [16]. CAF16 and CAF23 generated recently with retinoic acid (RA) as the immunomodulator, induce strong Ag-specific IgA responses in mice [17]. CAF16 is a CAF01 liposome, enriched by RA and stabilizing cholesterol, whereas CAF23 stabilized by polyethylene glycol (PEG), consists of both CAF16 and fast draining RA-enriched 1,2-distearoyl-*sn*-glycero-3-phosphocholine (DSPC) liposomes, having the ability to precondition the lymph node environment with RA, and generate lymphocytes with the potential to home in the mucosa [17].

Another important element to consider during vaccine development is selecting the most optimal route of administration. Different vaccine administration routes can lead to stimulation of different dendritic cell subsets presenting the Ags [3], which drain into different lymph node environments. For instance, induction of CD8^+^ T cell responses by CAF09 adjuvant (a previous version of CAF09b with a higher dose of Poly(I:C) immunomodulator) is dependent on i.p. administration, whereas intramuscular (i.m.) or subcutaneous administration elicit significantly lower CD8^+^ T cell responses [14], [18]. While i.m. immunization led to formation of a vaccine depot at the site of injection and poor self-drainage, i.p. immunization resulted in effective draining into the mediastinal and tracheobronchial lymph nodes [18]. Further evidence suggests that administering a vaccine into the *Houhai acupoint* in the dorsal midline between the tail and the rectum, induces significantly higher antibody levels, splenocyte proliferation and cytokine gene expression, compared to conventional immunization in the neck region of rats [19]. These recent findings clearly indicate that vaccine administration via an appropriate route is decisive for a desired immunological outcome. Here, the widely used i.m. injection was included as a standard administration route and compared with less traditional i.p. injection, included based on previously published data on CAF09 [18], and an unconventional perirectal (p.r.) injection into the ischiorectal fossa, a triangle-shaped space between the ischium and the rectum [20], with a presumable drainage into mucosa-associated lymphoid tissue around rectum.

The purpose of the present study was to use pigs to investigate immunomodulation of the adjuvants administered to different injection sites, in relation to other animal models. Successful translation into humans is highly dependent on choosing an appropriate animal model mimicking the human population well. Outbred large animal models, such as pigs, have numerous advantages, in particular because of the highly similar immune system and the natural susceptibility to some of the human pathogens. Vaccine testing in these species is more representative and reduces the risk of failure later in the process [21]. Moreover, the pig model has already proven to be useful in several aspects of vaccine development, in regards to testing and improving vaccine types, adjuvants, Ags, and delivery strategies [22].

In the current study, all adjuvants were formulated with a model Ag CTH522 protein from *Chlamydia trachomatis* (*C. trachomatis*). CTH522 Ag formulated with CAF01 is a human vaccine candidate which has completed phase I clinical trial for safety and immunogenicity [12]. Based on previously published data, we hypothesized that both adjuvant and the injection site would influence the immune response against the CTH522 Ag, by polarization towards different T cell subsets and generation of different antibody isotypes. More specifically, we hypothesized that CAF01 effect would be favored by the i.m. injection and give rise to a mixed population of Th1/Th17 cells; CAF09b induction of cytotoxic CD8^+^ T cells would be favored by the i.p. injection, and CAF16 and CAF23 enhancement of serum IgA and mucosal secretory IgA (SIgA) would be favored by the p.r. injection. To verify this hypothesis, state-of-the-art techniques were used to evaluate humoral responses by isotype-specific ELISA and cell-mediated immunity (CMI) by flow cytometry, cytokine release assays and microfluidic high-throughput qPCR. Here, we describe immunological effects of administering a model vaccine adjuvanted with distinct adjuvants into different injection sites using pigs from a commercial farm in non-sterile, real-life conditions. To our knowledge, this is the first study giving a detailed, comprehensive analysis of immune responses in swine following administration of a variety of adjuvants and injection sites. The use of these adjuvants in pigs, a large, highly valid animal model, is novel and generates knowledge that can be further exploited for future vaccine design.

## 2. Materials and methods

### 2.1. Pigs

For the initial and the main study presented here, a total of 72 Danish Landrace/Yorkshire/Duroc crossbred female piglets aged five weeks, were included. At the time of weaning at four weeks of age, all piglets were routinely vaccinated against Porcine circovirus type 2 (Circovac, Ceva, France). All animal procedures were performed in accordance with both national and international guidelines, and complied with the ARRIVE guidelines. The Danish Animal Experiments Inspectorate (Animal permit ID: 2020-15-0201-00493) and the institutional animal ethics committee approved all procedures.

### 2.2. Immunizations

Animals received 50 μg of purified *C. trachomatis* CTH522 Ag (Statens Serum Institut, Denmark; batch D-F0617-01) formulated in either CAF01, CAF09b, CAF16 or CAF23 adjuvant (Statens Serum Institut, Denmark). Each immunization comprised of 1 mL adjuvant mixed with 1 mL CTH522 Ag diluted in 10 mM Tris buffer + 2% glycerol, or just 10 mM Tris buffer + 2% glycerol for the non-adjuvanted control. Immunizations were delivered via either i.m., i.p. or p.r. route. For the initial experiment, all animals were immunized via p.r. route with CTH522 Ag formulated in CAF01, CAF16, CAF23 or without adjuvant, only in Tris buffer (Fig 1A). Each of the four groups included seven animals. In the main experiment, each of the nine groups included six animals, immunized via either i.m., i.p. or p.r. route with CTH522 Ag formulated in CAF01, CAF09b or CAF23 adjuvant (Fig 2A). Animals were primed (day 0) and subsequently boosted in two weeks’ interval (day 14), blood was drawn at day 0, 14, 21 and 28 post-prime immunization (p.i.). On day 28, animals were sacrificed and small intestinal samples were extracted for mucus collection (only for the main experiment) (Fig 1A and 2A).

**Fig 1:**
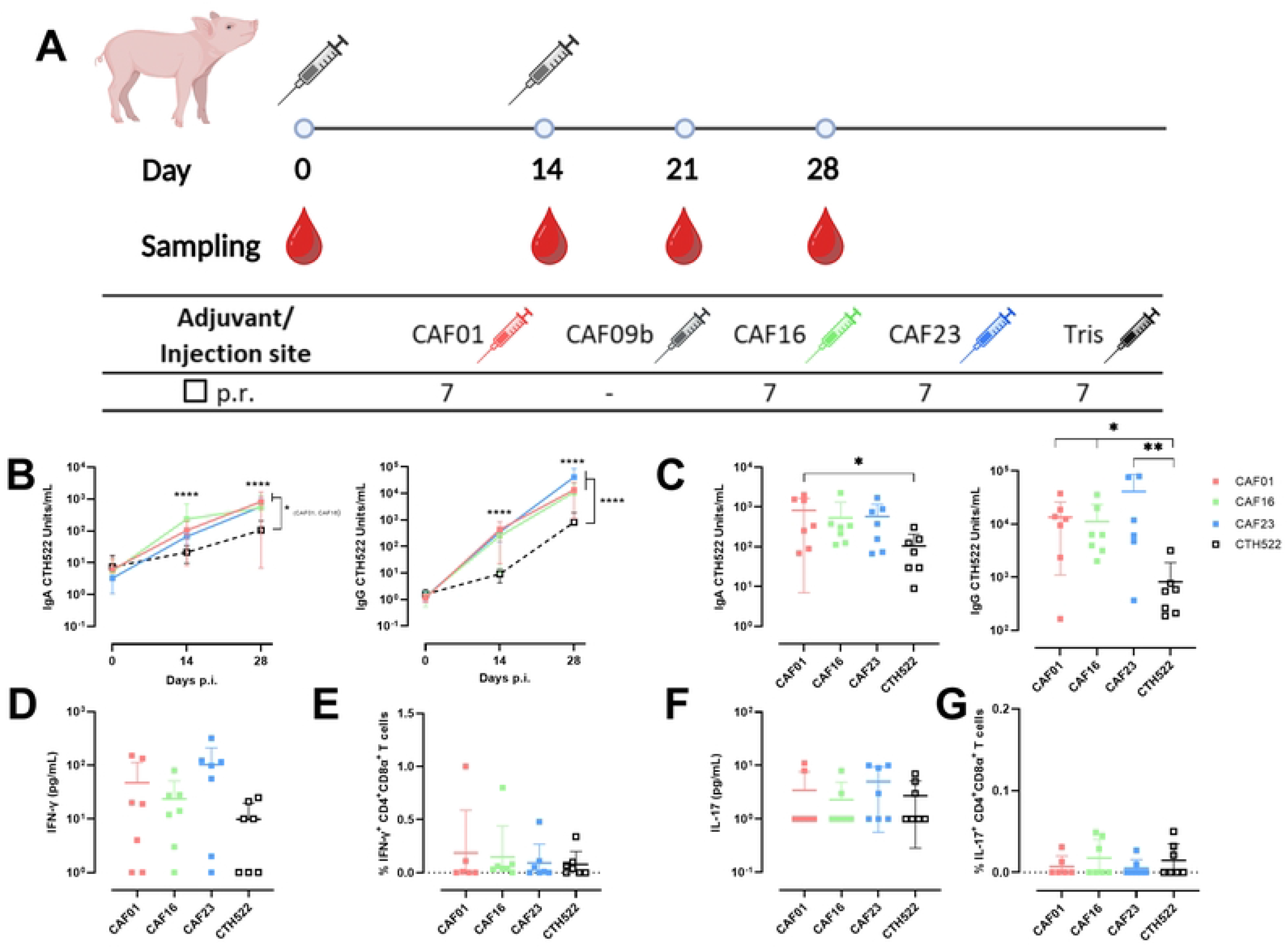
Initial experiment comparing three different CAFs formulated with CTH522 Ag versus non-adjuvanted CTH522 distributed via p.r. injection route: A) Experimental design – 28 outbred pigs were immunized via p.r. route with either CAF01:CTH522 (light pink), CAF16:CTH522 (green), CAF3:CTH522 (blue) or non-adjuvanted CTH522 (black) at day 0, and subsequently boosted at day 14. Blood for serum and/or PBMC isolation was drawn at day 0, 14, 21 and at final day 28. B) Serum IgA and IgG CTH522-specific responses to vaccination. C) Individual anti-CTH522 IgA and IgG responses at day 28. D) IFN-γ and F) IL-17 cytokine release assays of PBMCs re-stimulated with CTH522. The background from the NS controls was subtracted, and all animals were assigned a minimal IFN-γ and IL-17 concentration of 1 pg/mL. Percentage of E) IFN-γ-producing Th1 cells, and G) IL-17-producing Th17 cells as a proportion of CD4^+^CD8α^+^ T cells measured by ICS. The background from NS controls was subtracted, the pigs with higher background than the signal after CTH22 antigen re-stimulation were set to 0. B) – G) Data represent one experiment, and all pigs (28) were initially included. All data were log-transformed prior to statistical analysis. Statistically significant differences are indicated by *, **, *** or **** where p < 0.05, p < 0.01, p < 0.001 or p < 0.0001 respectively (one-way ANOVA, followed by Dunnett’s multiple comparisons test). There is no statistically significant difference, unless otherwise indicated. Data are presented as mean ± SD with data points representing responses from individual animals.

**Fig 2:**
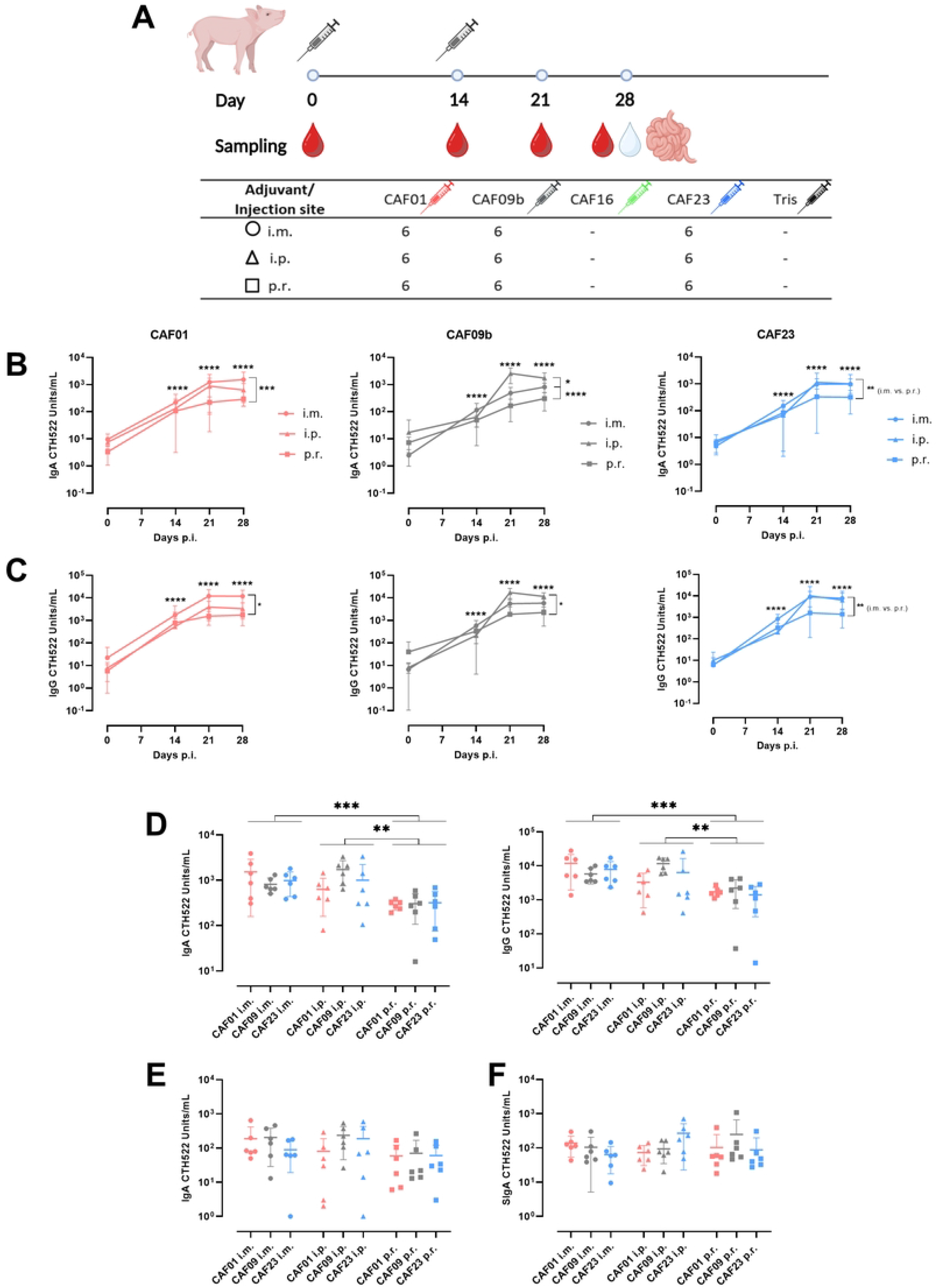
CTH522-specific antibodies measured by ELISA after prime and boost immunization comparing three different adjuvants and three different injection sites: A) Experimental design – 54 outbred pigs were immunized via i.m. (circle), intraperitoneal i.p. (triangle) or p.r. (square) injection route with either CAF01:CTH522 (light pink), CAF09b:CTH522 (grey) or CAF23:CTH522 (blue) at day 0, and subsequently boosted at day 14. Blood for serum and/or PBMCs was drawn at day 0, 14, 21 and 28; and the mucus from the small intestine was isolated at the final day 28. Detection of CTH522-specific serum B) IgA (top row) and C) IgG (bottom row) comparing i.m., i.p. or p.r. site of injection across all time points. Each adjuvant group is shown separately. D) Comparison of the anti-CTH522 IgA and IgG antibodies, respectively, in serum across all groups at day 28. Comparison of CTH522-specific E) IgA and F) SIgA antibodies, in small intestinal mucus across all groups at day 28. B) – F) Data represent two independent experiments, and all pigs (54) were included. All data were log-transformed prior to statistical analysis. Statistically significant differences are indicated by *, **, *** or **** where p < 0.05, p < 0.01, p < 0.001 or p < 0.0001, respectively (two-way ANOVA followed by Tukey’s multiple comparisons test). There is no statistically significant difference, unless otherwise indicated. Data are presented as mean ± SD with data points representing responses from individual animals.

### 2.3. Cell isolation

Peripheral blood samples were collected from the jugular vein into BD Vacutainer EDTA tubes (BD Diagnostics, catalogue number (cat.): 366643). Peripheral blood mononuclear cells (PBMCs) were purified using SepMate™ tubes (StemCell Technologies, cat.: 85450) according to the manufacturer’s protocol. Briefly, EDTA blood was diluted 1:1 in sterile phosphate buffered saline (PBS), carefully pipetted into SepMate™ tubes with density gradient medium Lymphoprep (StemCell Technologies, cat.: 07851) and separated by centrifugation for 20 min, 1200 G at room temperature (RT). The cells were washed twice in PBS + 2% fetal calf serum (FCS; Biowest, cat.: S181B-500), followed by red blood cells lysis. PBMCs were resuspended in freezing medium consisting of FCS/40% RPMI 1640 (Thermo Fisher Scientific, cat.: 11875093) + 10% dimethyl sulfoxide (DMSO), and a total of 1×10^7^ PBMCs per vial were cryopreserved for subsequent analyses. Cryotubes with cells in freezing medium were placed into CoolCell LX freezing containers (BioCision, USA) at -80°C, and transferred into liquid nitrogen after 24 h for long-term storage.

### 2.4. Mucus isolation

Small intestinal mucus was isolated to monitor mucosal immune responses. Pigs were sedated with Zoletil mix i.m. (12.5 mg tiletamine, 12.5 mg zolazepam, 12.5 mg xylazine, 12.5 mg ketamine per mL at 1 mL/15 kg), euthanized by captive bolt pistol and bled. Two 8 cm segments of small intestine at 50 cm and 150 cm distal to the duodenojejunal flexure were isolated, filled with 5 mL PBS/0.01 M EDTA/protease inhibitor cocktail buffer (Sigma-Aldrich, cat.: P8340) and transferred to a cold box. Afterwards, the intestinal samples were cut open and the mucus layer was gently removed and centrifuged for 20 min, 4100 G. The supernatants were collected and stored at -20°C. To inhibit non-specific SIgA binding in mucus, 1% Tween 20 was added to the thawed samples for 30 min at RT. The samples were centrifuged and supernatants were collected and stored at -20°C until analysis.

### 2.5. Antibody ELISA

The Ag-specific antibodies in serum and small intestinal mucus were monitored by indirect ELISA. Nunc MaxiSorp 96-well plates (Thermo Fisher Scientific, cat.: 442404) for the analysis of serum samples, and Nunc PolySorp 96-well plates (Thermo Fisher Scientific, cat.: 475094) for mucus samples, were coated with 1 μg/mL vaccine Ag CTH522 in carbonate buffer (SSI Diagnostica) overnight at 4°C. IgA and IgG antibody isotypes were detected by horse reddish peroxidase (HRP)-conjugated goat anti-pig antibodies specific for porcine IgA (clone AAI40P, BioRad) diluted 1:5 000; and porcine IgG (clone AAI41P, BioRad) diluted 1:20 000. SIgA was detected by HRP-conjugated streptavidin using biotinylated mouse anti-pig secretory component (clone K60 1F1, BioRad) diluted 1:2 000. The reactions were developed with 50 μL/well of tetramethylbenzidine substrate (TMB; Kem-En-Tec, cat.: 4395A) and stopped with 0.5 M sulfuric acid. All samples were included in technical duplicates. The plates were read on a microplate reader (TECAN Sunrise) at 450 nm/620 nm. For mucus ELISA, a pool of mucus samples was used as standard, while for serum ELISA, a commercial pig serum (PS; Biowest, cat.: S2400) was used as a standard, because of a natural low cross-reactivity to CTH522, presumably due to *Chlamydia suis* (*C. suis*). Serially diluted standards were assigned arbitrary anti-CTH522 IgA and IgG units with a value of 100 units/mL. Anti-CTH522 units for each sample were calculated by log-log transformation against the standard. All samples were given a minimum value of 1 unit/mL.

### 2.6. Cytokine ELISA

IFN-γ and IL-17A concentrations in the cell culture supernatant were determined by ELISA. Cryopreserved PBMCs were thawed and cell counts were determined by Nucleocounter NC-200 (Chemometec, Denmark). A total of 1×10^6^ viable cells per well were rested for 7 h in RPMI 1640 + 10% PS in the incubator (37°C, 5% CO_2_). For IFN-γ ELISA, the cells were re-stimulated in technical triplicates with 5 μg/mL CTH522, or 1 μg/mL Staphylococcal enterotoxin B (SEB; Sigma Aldrich, cat.: S4881) as positive control, or media alone as negative control. The cell culture supernatant was collected after 18 h of incubation, while the cells were preserved in RNAlater (Ambion, cat.: AM7021) with 10% PBS and frozen at -20°C for subsequent RNA isolation described below. MaxiSorp plates were coated with 1 μg/mL of monoclonal mouse anti-pig IFN-γ antibody (clone P2F6, Thermo Fisher Scientific) in carbonate buffer, or polyclonal rabbit anti-pig IL-17A (clone KP0498S-100, Kingfisher Biotech) in PBS, and incubated overnight at 4°C. A biotinylated mouse-anti pig antibody (clone P2C11, Thermo Fisher Scientific) diluted 1:500 was used for detection, followed by HRP-conjugated streptavidin (Thermo Fisher Scientific, cat.: SNN1004) diluted 1:10 000. For IL-17A ELISA, the cells were stimulated in the same manner, but the cell culture supernatant was collected after 72 h, and the cells were not kept for further analysis. A biotinylated rabbit anti-pig antibody (clone KPB0499S-050, Kingfisher Biotech) at 100 ng/mL was used for detection, followed by HRP-conjugated streptavidin diluted 1:10 000. The plates were developed with 50 μL/well of TMB substrate and the reaction was stopped with 0.5 M sulfuric acid. Each plate included a duplicate of standard with a known cytokine concentration. In-house IFN-γ standard was titrated in a serial 2-fold dilution, and IL-17A standard (clone RP0128S-005, Kingfisher Biotech) was prepared following manufacturer’s instructions and used for making a log-transformed standard curve. The lowest point of the standard curve used for calculations of the cytokine concentrations was 6 pg/mL, therefore, all the calculated concentrations below 6 pg/mL were set to 6 pg/mL. The background cytokine production measured in NS control was subtracted from the respective CTH522 re-stimulated sample, and calculated values lower than 1 pg/mL were set to 1 pg/mL.

### 2.7. IFN-γ ELISpot

IFN-γ production was evaluated by ELISpot assay, as described in [23], following stimulation with CTH522 vaccine Ag and a CTH522 peptide pool consisting of twelve 9-mer and 10-mer peptides. Pigs were typed for the most common swine leukocyte antigen (SLA) alleles carried by Danish pigs [24], allowing us to use the NetMHCPan 4.1 tool to predict immunogenic 9-mer and 10-mer peptides from the CTH522 protein. The best candidate peptides were custom-synthesized (Schafer-N, Denmark). Briefly, MultiScreen® IP Filter Plates with hydrophobic polyvinylidene fluoride membrane (Merck Milipore, cat.: MAIPS4510) were pre-wet with 35% ethanol in sterile MiliQ water for 1 min and coated with 5 μg/mL of monoclonal mouse anti-pig IFN-γ antibody (clone P2F6, Thermo Fisher Scientific) overnight at 4°C. The plates were blocked for a minimum of 1 h at 37°C in serum-free AIM-V™ medium (Thermo Fisher Scientific, cat.: 12055091). To each well, a total of 5×10^5^ thawed PBMCs were added and incubated for 20 h at 37°C, in the presence of 5 μg/mL CTH522, the CTH522 peptide pool at final concentration of 2 μM each, 1 μg/mL SEB for the positive control, or media alone for the negative control. Biotinylated mouse anti-pig IFN-γ antibody (clone P2C11, Thermo Fisher Scientific) for detection was added at 1 μg/mL and incubated for 1 h at RT. Streptavidin alkaline phosphatase conjugate (Promega, cat.: V5591) was diluted 1:2 500 and incubated for 1 h at RT. Finally, 5-bromo-4-chloro-3-indolyl phosphate (BCIP)/nitro blue tetrazolium (NBT) substrate (Thermo Fisher Scientific, cat.: 34042) was added and the reaction was terminated after 5 min under running tap water. The plates were let to air-dry overnight in the dark and were read on ImmunoSpot® Analyzer S6 Ultra (Cellular Technology Limited, USA), using the SmartCount feature with manual quality control. The data is presented as number of spot-forming cells (SFCs) per 5×10^5^ cells, calculated as an average of the three technical repeats, shown after subtraction of the SFCs from the NS media control.

### 2.8. Flow cytometry

Cryopreserved PBMCs were thawed and cell counts were determined by Nucleocounter NC-200. A total of 1 – 2×10^6^ viable cells per well were rested for 7 h in RPMI 1640 + 10% PS in the incubator (37°C, 5% CO_2_). Afterwards, the cells were re-stimulated with 5 μg/mL CTH522 or 1 μg/mL phytohemagglutinin (PHA; Roche, cat.: 11249738001) as positive control, or RPMI 1640 + 10% PS as negative non-stimulated (NS) control for 15 h followed by additional 4 h in the presence of 5 μg/mL Brefeldin A (Sigma-Aldrich, cat.: B7651-5MG) and 2 μM monensin (Thermo Fisher Scientific, cat.: 00-4505-51). Cells were surface-stained for 30 min at 4°C. Cytofix/Cytoperm™ Fixation/Permeabilization Solution Kit (BD Biosciences, cat.: 554714) was used according to manufacturer’s instructions, followed by intracellular cytokine staining (ICS) for 30 min at 4°C. In case of FoxP3 detection, cells were treated with FoxP3 Transcription Factor Fixation/Permeabilization Kit (Thermo Fisher Scientific, cat.: 00-5521-00) according to manufacturer’s instructions, followed by incubation with FoxP3 antibody for 30 min at 4°C. All antibodies were titrated and used at pre-determined optimal concentrations (S1 Table). Flow cytometry measurements were performed using LSRFortessa™ Cell Analyzer (BD Biosciences, USA) equipped with three lasers (405, 488 and 640 nm) and a high throughput sampler. Compensation was calculated automatically using single-stained controls prior to each measurement. The data was analyzed by FlowJo Data Analysis Software version 10.8.1. Fluorescence minus one (FMO) controls were used to set cytokine gates, and gating strategies for data analysis are shown in S1 Fig. The background cytokine production in the NS control was always subtracted from the CTH522 re-stimulated samples to give the Ag-specific signal only.

### 2.9. RNA extraction and primer design

Total RNA was isolated from 2×10^6^ PBMCs preserved in RNALater + 10% PBS. PBMCs were stimulated as described in section 2.6. RNA was isolated using RNeasy Mini Kit (Qiagen, cat.: 74104), according to manufacturer’s instructions, and included DNA digestion using RNAse-free DNAse kit (Qiagen, cat.: 79254). Total RNA was quantified by UV spectrometry at 260 nm and RNA purity was estimated by 260/230 and 260/280 absorbance ratios using a NanoDrop 2000 spectrophotometer (Thermo Fisher Scientific, USA). In addition, RNA integrity was evaluated by Agilent® 2100 Bioanalyzer (Agilent Technologies, Germany), using Agilent RNA 6000 Nano Kit (Agilent Technologies, cat.: 5067-1511), according to manufacturer’s instructions. The mean RIN value of 9 ± 0.5 (interval 7.6 – 10), demonstrating integrous, high-quality input RNA across the samples. The eluted RNA in RNAse-free water was stored at -80°C.

Primers were designed based on sequences from *Sus scrofa,* published on public databases GenBank (NCBI) and Ensembl using Primer3Plus version 4.1.0. tool (https://www.primer3plus.com/), and custom-synthesized (Eurofins, Germany).

### 2.10. Microfluidic high-throughput qPCR

As the porcine immunological toolbox is still limited compared to small rodents and human, several markers we wished to monitor are not available as specific anti-porcine monoclonal antibodies. Therefore, microfluidic high-throughput qPCR was performed as described in detail in [25]. Briefly, complementary DNA (cDNA) from each RNA sample was synthesized in two separate technical replicates from 150 ng of total RNA using QuantiTect Reverse Transcription Kit (Qiagen, cat.: 205311), according to manufacturer’s instructions. All samples were diluted ten times in low EDTA TE-buffer (VWR, cat.: APLIA8569.0500) in a total volume of 20 μL. A preamplification primer mix consisting of 20 μM of each primer pair was mixed with 5 μL TaqMan PreAmp Master Mix (Applied Biosystems, cat.: PN 4391128) and added to each of the cDNA samples. The mix was incubated 10 min at 95°C followed by 18 cycles of 15 s at 95°C and 15 s at 60°C. Afterwards, 4 U/μL Exonucluease I (New England Biolabs, cat.: PN MO293L) was added to the preamplification mix and incubated in the thermocycler for 30 min at 37°C and 15 min at 80°C to remove non-incorporated primers.

To choose the most suitable gene targets and the best-performing primers for the genes of interest, an initial qPCR screening of 39 putative primer assays in a 48.48 Dynamic Array (48 cDNA samples and 48 gene targets) using the BioMark thermocycler (Fluidigm Corporation, USA) was performed. The 48.48 Dynamic Array with loaded control line fluid was primed in IFC Controller MX (Fluidigm Corporation, USA). Assay mastermix was prepared from 2× Dynamic Array Sample & Assay Loading Reagent Kit (Fluidigm Corporation, cat.: 85000825) and 1× low EDTA TE-buffer and mixed with 20 μM of a specific primer pair. Pre-sample mix was prepared from 2× TaqMan Gene Expression Master mix (Applied Biosystems, PN 4369016), 20× DNA binding dye, 20× EvaGreen (VWR, cat.; BTIU31000) and 1× low EDTA TE-buffer. Finally, 1.5 μL of each preamplified cDNA was distributed. Mastermix and samples were loaded on the 48.48 Dynamic Array, which was placed back into the IFC Controller, and finally into BioMark instrument. The thermal protocol was as following: 2 min at 50°C, 10 min at 90°C; qPCR consisted of 35 cycles of 15 s at 95°C (denaturation) and 60 s at 60°C (annealing and extension), followed by a melt curve program of 30 s at 60°C and 1°C/3 s until 95°C.

The final primer set (sequences listed in the S2 Table), consisting of 20 genes and four reference genes, and all samples were loaded on two 192.24 Dynamic Arrays (192 cDNA samples and 24 gene targets). The 192.24 Dynamic Array was prepared as described above, but was primed only once in RX IFC Controller (Fluidigm Corporation, USA). The thermal protocol was as following: 2 min at 50°C, 30 min at 70°C and 10 min at 25°C, followed by 2 min at 50°C and hot start for 10 min at 95°C. The rest of the qPCR cycle was identical with the one described above for 48.48 Dynamic Array. Controls without reverse transcriptase and non-template controls were included to monitor any DNA contamination and non-specific amplification. Interplate calibrators were included to correct for any plate-to-plate variation.

The data quality was assessed using Fluidigm Real-Time PCR Analysis software v.4.8.1 (Fluidigm Corporation, USA) and demonstrated outstanding quality proven by the high reproducibility between cDNA replicates. Data pre-processing and analysis were done using GenEx v.7 (MultiD, Sweden), as described in great detail in [25]. The steps included the following: 1. Visual evaluation of amplification and melting curves, 2. Correction of PCR amplification efficiency using calibration curves from dilution series, 3. Interplate calibration, 4. Normalization to reference genes – four reference genes (*ACTB, HPRT, RPL13A* and *YWHAE*) were validated by geNorm and NormFinder algorithms incorporated in GenEx v.7, 5. Calculation of averages from cDNA technical replicates, 6. Calculation of relative expression using the NS control group as a baseline, 7. Log_2_ transformation: For all statistical analyses, Log_2_-transformed data corresponding to relative quantity (RQ) were used, and the fold change was calculated as a ratio of RQ of the CTH522-stimulated sample and the RQ of the respective NS sample. For heatmap data presentation, the Log_2_ RQ values of gene expression were transformed into Z-score to normalize the differences between the genes.

### 2.11. Statistical analysis

Statistical comparisons were performed either via ordinary one-way ANOVA, or ordinary two-way ANOVA, followed by Tukey’s multiple comparisons test or Dunnett’s multiple comparisons test. As significant, p < 0.05 (*) was considered, and p < 0.01 (**), p < 0.001 (***) and p < 0.0001 (****) are indicated. There was no significant difference, if not otherwise indicated. Data are shown as mean ± standard deviation (SD). GraphPad Prism Version 9.3.1 for Windows (California, United States) was used for all statistical analyses.

## 3. Results

### 3.1. The CAF-adjuvanted vaccines are immunogenic in pigs

CAF adjuvants have been developed to induce strong CMI, but induce humoral responses too [26], allowing us to use the antibody responses as an easily-accessible proof of vaccine immunogenicity in the initial experiment (Fig 1). Using indirect ELISA, we evaluated whether the amount of Ag-specific IgA and IgG antibodies in serum were affected by the adjuvant. All animals were serologically naïve to CTH522 prior to the immunization (Fig 1B).

CTH522 was shown to be immunogenic also on its own, as the anti-CTH522 IgA and IgG antibodies in the non-adjuvanted group increased 10 and 1000-fold respectively, between day 0 and day 28, but remained lower than in all adjuvanted groups. Overall, IgA antibody units induced by the non-adjuvanted CTH522 Ag were on average 6-fold lower than in all adjuvanted groups (CAF01 and CAF16 vs. CTH522 p = 0.0244 and p = 0.0409, respectively) (Fig 1B). CAF23 had no significant effect on IgA in serum overall (p = 0.0969), compared to the non-adjuvanted control. In case of anti-CTH522 IgG antibodies, the effect of all adjuvants over time was more pronounced than for IgA (p < 0.0001), giving 13-fold higher number of antibody units in the CAF16 group, and rising up to 50-fold in the CAF23 group, compared to non-adjuvanted CTH522 (Fig 1B). Adjuvant effect on the antibody level can be seen already at day 14, two weeks after the prime immunization, and was enhanced by the booster immunization (Fig 1B). When only the final day 28 is considered, there was a significant difference in IgA antibody units between the non-adjuvanted group and CAF01 group (p = 0.0198), while CAF16 and CAF23 groups were borderline significant (p = 0.0569 and p = 0.0518, respectively). Serum IgG was significantly increased in CAF23 group (p = 0.0015) and also CAF01 and CAF16 groups (p = 0.0136 and p = 0.0112, respectively) as compared to the non-adjuvanted control (Fig 1C).

### 3.2. Cell-mediated immune responses were not polarized by adjuvant immunomodulators

We monitored IFN-γ and IL-17, as CAF01 was shown to induce a strong mixed Th1/Th17 immune response in mice and non-human primates [5], [27]. IFN-γ concentration was variable within each group, but there was a tendency towards CAF23 group producing the highest amounts of IFN-γ, even though two pigs were non-responders (Fig 1D). IL-17 production was low and after background subtraction did not exceed 12 pg/mL, with no differences between groups (Fig 1E). The SEB-stimulated positive controls and the NS controls are shown in S3 Fig.

The current consensus on the phenotype of porcine Th cells is represented by the CD3^+^TCRγδ^-^ CD4^+^CD8α^+/-^ population [28], and based on the cytokine production these can be subdivided into Th1 cells producing IFN-γ and Th17 cells producing IL-17. The gating strategy used for data analysis is shown in S1B Fig. Pigs with negative Ag-specific signal were regarded as non-responders and their Ag-specific responses were set to 0. There were no clear differences in cytokine production in the CD4^+^CD8α^+^ effector population (Fig 1F-G). The low frequency of IL-17^+^ T cells may be partly attributed to the use of a sub-optimal cross-reactive human clone, published also elsewhere [29], as porcine-specific antibody for flow cytometry is currently not available commercially. The cytokine production in PHA-stimulated positive controls and NS controls are shown in S3C-D Fig.

### 3.3. The effect of injection site was greater than the effect of the adjuvant for antibody responses

In the next experiment, CAF01, CAF09b and CAF23 adjuvants were compared using different routes of administration, allowing us to examine both the influence of adjuvants and administration routes on the resolving immune responses (Fig 2A).

All pigs were serologically naïve to CTH522 prior to the vaccination on day 0 (Fig 2B). Both IgA and IgG antibodies were increased in the serum already at day 14, two weeks after the prime immunization in all adjuvant groups (Fig 2B-C). Following the boost, antibody responses were further enhanced at day 21, when significant differences between the injection sites were clear. The same trend with comparable antibody concentrations was observed at day 28 (Fig 2B-C), when the antibody levels reached a plateau. Overall, in the CAF01 group, i.m. injection induced significantly more IgA and IgG antibodies, than p.r. injection (p = 0.0007 and p = 0.0109, respectively) and likewise in the CAF23 group (p = 0.0061 and p = 0.0007, respectively). In the CAF09b group, i.p. injection induced significantly more IgA than both i.m. (p = 0.0342) and p.r. (p < 0.0001) injections, while in case of IgG, only i.p. injection induced significantly higher antibody units than the p.r. injection (p = 0.0140). Comparing the antibody levels at day 28, IgA serum units were significantly higher in i.m. and i.p. immunized animals, compared to the p.r. injected counterpart (p = 0.0002 and p = 0.0013, respectively). Likewise, for IgG antibodies, both i.m. and i.p. immunization gave significantly higher responses than the p.r. route (p = 0.0002 and p = 0.0066, respectively). Administering the vaccine through both i.m. and i.p. route induced significantly higher anti-CTH522 IgA and IgG antibodies in serum (Fig 2D). There was no significant difference between the adjuvant groups at any time point.

### 3.4. CAF23 did not induce significantly higher mucosal IgA and SIgA response

Since the RA-containing CAF23 adjuvant was shown to drive significant serum and mucosal IgA responses after parental vaccination in mice [17], we monitored IgA and SIgA – the principal effective form of mucosal IgA [30], in the small intestinal mucus. Unlike in mice, IgA levels in mucus were not increased in CAF23-immunized animals when compared to CAF01 and CAF09b, but rather the levels were evenly distributed between the groups (Fig 2E), following the adjuvant trend from serum IgA (Fig 2D). There were no significant differences in mucus IgA levels between the injection routes, in contrast to the differences observed in serum (Fig 2D). Furthermore, SIgA levels in mucus were not significantly different between the adjuvant groups, but in this case, the CAF23 i.p. group gave overall the highest SIgA units on average (Fig 2F), which aligned with our original hypothesis. The p.r. injection was hypothesized to activate mucosa-associated lymphoid tissue around rectum and subsequently induce mucosal response, but it induced similar IgA and SIgA antibody levels as i.m. and i.p. injections. It is notable that in mice, neither serum nor mucosal IgA is induced by CAF01 when administered subcutaneously [31], or i.m., while in pigs, both antibodies were present in serum and mucus, regardless of the adjuvant.

### 3.5. Gene expression analysis resulted in clustering of samples based on stimulation rather than type of adjuvant or injection site

The overall relative gene quantities (Log_2_ RQs) are presented by the respective adjuvant and stimulation, and as individual animals sorted in groups based on the stimulation, adjuvant and injection site (Fig 3). As an addition to flow cytometry staining (Fig 4), genes specifying different T cell subsets (*IL2, IL10, IL17A, IFNG, TGFB, TNFA, FOXP3, RORC;* Fig 4), T cell activation (*KLRK1, GZMB, EOMES, LAMP1;* Fig 4 and S6 Fig), and homing (*CCR6, CCR7, ITGA4, ITGB7;* Fig 6) as well as central genes from the pathways activated by CAF01 (*CLEC4E, CARD9;* S6 Fig) and CAF09b (*STAT1, DDX59;* S6 Fig) adjuvants in mice, were chosen for primer design and subsequent analysis. Based on the readouts from the screening 48.48 Dynamic Array, we narrowed down the genes of interest to above-stated 24 genes, including four validated reference genes. One limitation of our qPCR experiment was the lack of a control group without immunization or adjuvant. Therefore, as a control, NS samples have been used, similarly like for flow cytometry, cytokine ELISA and ELISpot assays performed here. Relative expressions of the genes are shown as Log_2_ RQ, and the fold changes were calculated as a ratio of CTH522 re-stimulated and NS control Log_2_ RQs (Fig 4).

**Fig 3:**
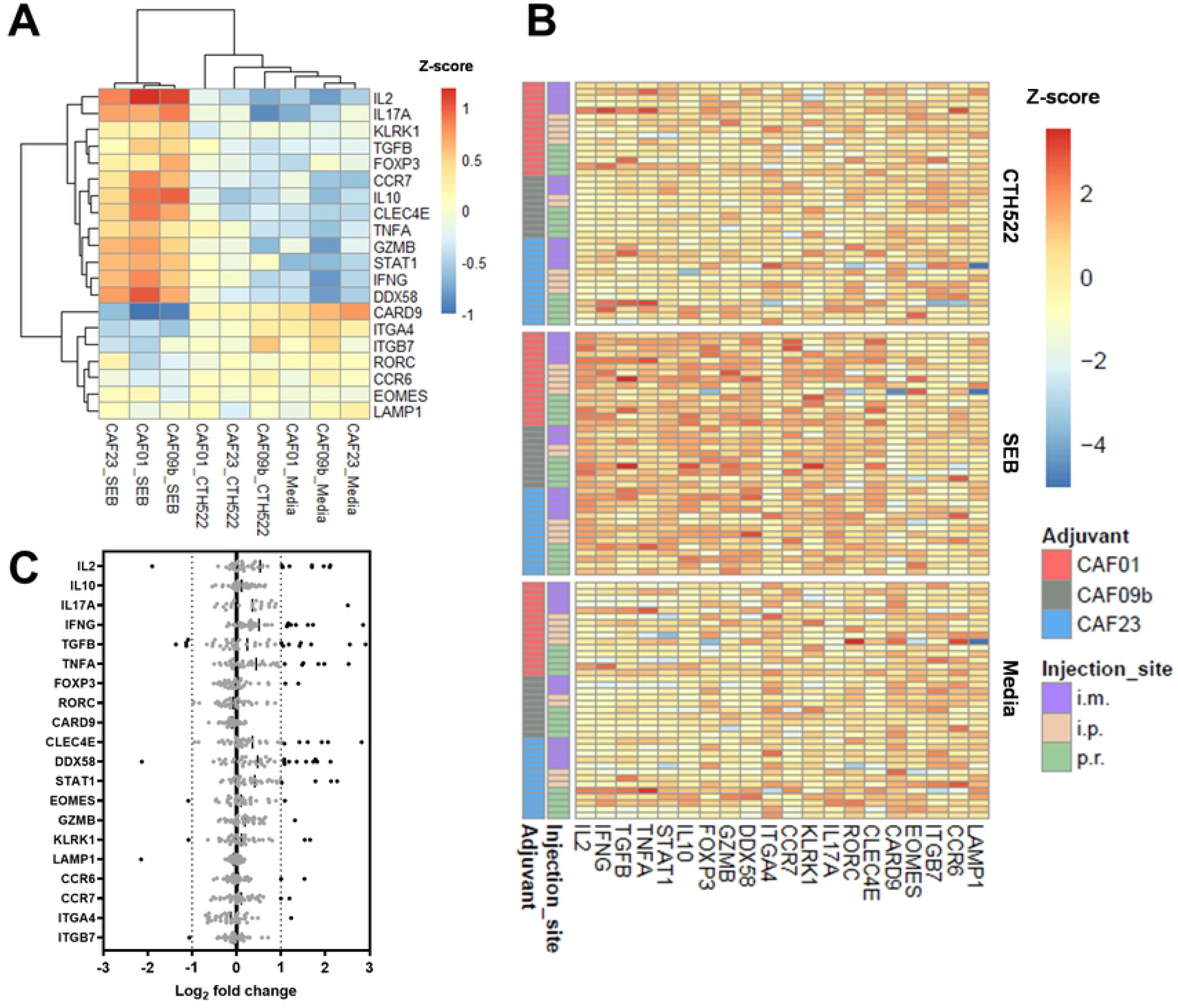
High-throughput qPCR: A) Heatmap of an average relative gene expression of 20 genes. The samples were grouped by the adjuvant and stimulation (SEB, CTH522 and media) and clustered based on correlation. B) Relative gene expression heatmap of individual pigs divided into three groups based on re-stimulations, sorted by adjuvant and injection site, as indicated. Both heatmaps are represented by the Z-score. C) Scatterplot of 20 genes depicting expression fold changes. Relative gene quantity (RQ) of CTH522 Ag re-stimulated PBMCs was divided by the respective RQ of the NS control and referred to as Log_2_ fold change. The solid black line represents Log_2_ 0-fold change, and the dotted lines represent Log_2_ 1-fold downregulation and 1-fold upregulation, respectively. Points represent relative fold change in individual pigs – black points denote fold changes > 1-fold up- or downregulation, while grey points denote no significant change. Mean fold change for each gene is displayed as a black line. One outlier from the CAF09b i.p. group was excluded, due to low reproducibility between the cDNA replicates.

**Fig 4:**
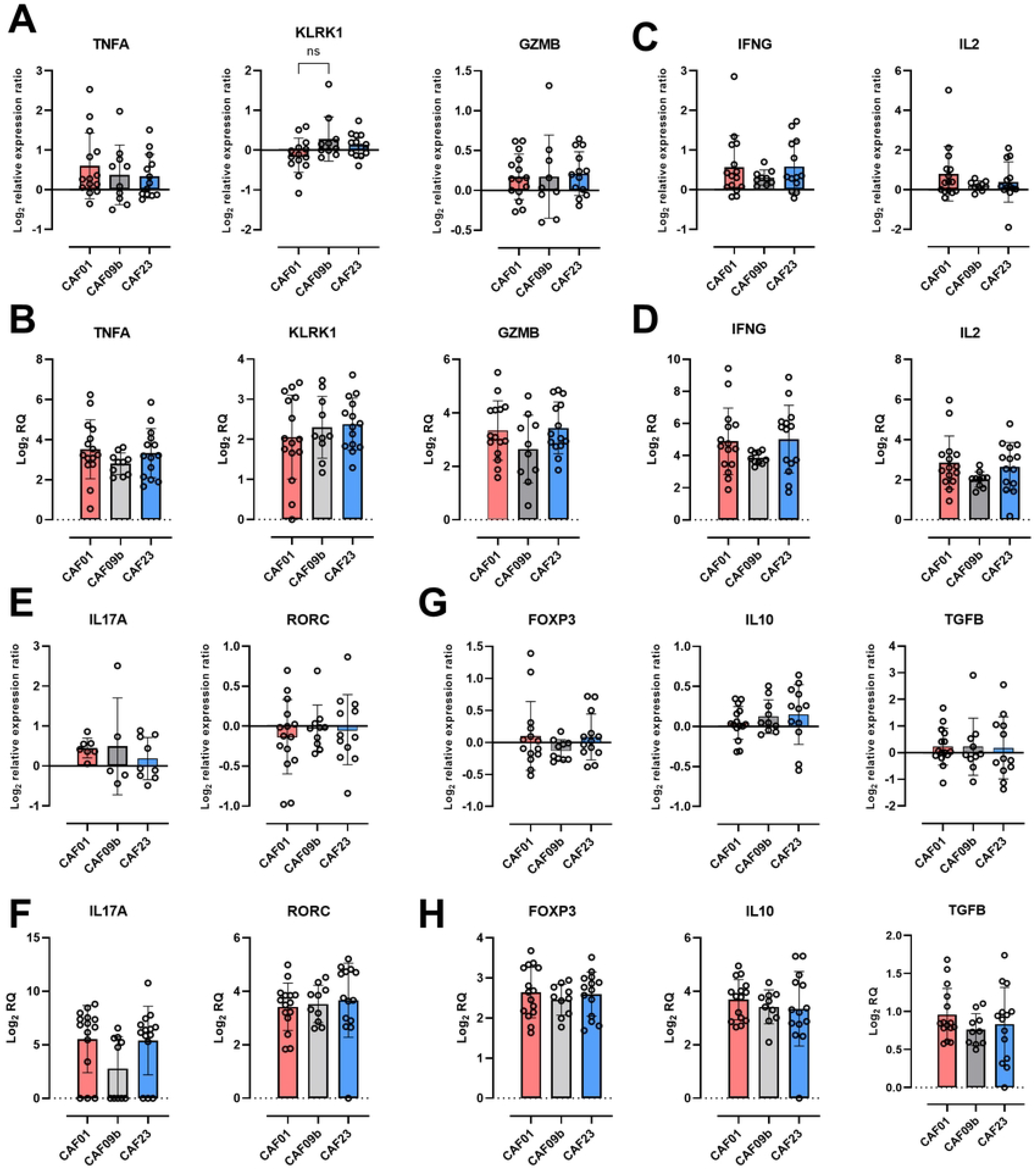
Expression of genes characterizing different T cell subsets relevant for cell-mediated immune responses measured by high-throughput qPCR: A) CTL population: *TNFA*, *KLRK1* and *GZMB* Log_2_ fold change, and B) Log_2_ RQ in CTH522-re-stimulated PBMCs. C) Th1 population: *IFNG* and *IL2* Log_2_ fold change, and D) Log_2_ RQ in CTH522-re-stimulated PBMCs. E) Th17 population: *IL17A* and *RORC* Log_2_ fold change, and F) Log_2_ RQ in CTH522-re-stimulated PBMCs. G) Treg population: *FOXP3*, *IL10* and *TGFB* Log_2_ fold change, and H) Log_2_ RQ in CTH522-re-stimulated PBMCs. A) – H) Fold changes are represented as a ratio between relative gene quantity in CTH522 re-stimulated PBMCs and NS PBMCs. Data represent two independent experiments, and 40 randomly chosen pigs were initially included. One outlier from the CAF09b i.p. group was excluded, due to low reproducibility between the cDNA replicates. The groups are based on adjuvants only, the injection sites were pooled under the respective adjuvant for the analysis. No statistically significant differences were identified by one-way ANOVA. Data are presented as mean ± SD with data points representing responses from individual animals.

In Fig 3A, heatmap clustering was performed based on correlation, and clearly distinguished gene expressions between the different stimulations, especially by separating SEB-stimulated samples in one big cluster, and CTH522-stimulated and NS samples in another cluster. The latter cluster subsequently sub-clustered into a separate CAF01-CTH522 cluster and a mixed CTH522/NS (media) cluster. This indicates that CAF01 group re-stimulated with CTH522 had the largest effect on differential gene expression, while the rest of the CTH522 groups were closely resembling the NS groups. In Fig 3B, the heatmap presenting all individual animals, confirms the high variability between the animals, and depicts the upregulation of several genes in SEB-stimulated samples, but no differences between NS and CTH522 re-stimulated animals, or between the adjuvants and injection sites. Log_2_ fold changes (a ratio between CTH522 and NS RQs) in all the animals for the 20 investigated genes are summarized in Fig 3C. The cut-off fold changes were set to Log_2_= ±1, as a significant down- or up-regulation. Thereafter, all the genes were analyzed separately and the relative expression as well as fold changes between CTH522 and NS were pooled under the respective adjuvant or injection site and are shown in Figs 4, 6 and 7 and S6 Fig. The individual expressions and fold changes are presented and discussed in the following section 3.6.

### 3.6. The type of adjuvant and injection site showed little polarizing effect on the cell-mediated immune responses

Gating strategies for the three antibody panels used for data analysis are presented in S1 Fig. Here, the cytokine signal after subtraction of the signal in the respective NS control is shown (Fig 5), positive control and NS control is shown in S4 Fig. In a non-negligible number of pigs, the background cytokine production was higher than in the CTH522 re-stimulated samples, and therefore these pigs were regarded as non-responders, and their Ag-specific responses were set to 0. The current understanding is that CD8α is expressed on activated and memory porcine CD4^+^ T cells [28]. Porcine cytotoxic T lymphocytes (CTLs) are represented by the CD3^+^TCRγδ^-^ CD4^-^CD8^high^αβ population [28], [33]. We monitored activated CTLs, which we defined as producers of IFN-γ and/or TNF-α and/or perforin (Fig 5A). The percentages represent a union of the aforementioned three cytokine gates (“Make Or Gate” function, FlowJo). The percentages are overall high due to high proportion of porcine T cells producing perforin. Since several pigs were non-responders, it is hard to be conclusive, but the tendency seemed to be towards CAF01 groups giving the highest cytokine proportions in the CTL population. In contrast with mouse studies [18], the CAF09b i.p. group had the lowest percentage of activated CTLs and four pigs did not respond to the Ag. In S2A Fig, representative flow cytometry dot plots of one animal from each group are displayed, showing IFN-γ^+^ T cells within the CD8α^+^CD8β^+^ CTL population after CTH522 re-stimulation.

**Fig 5:**
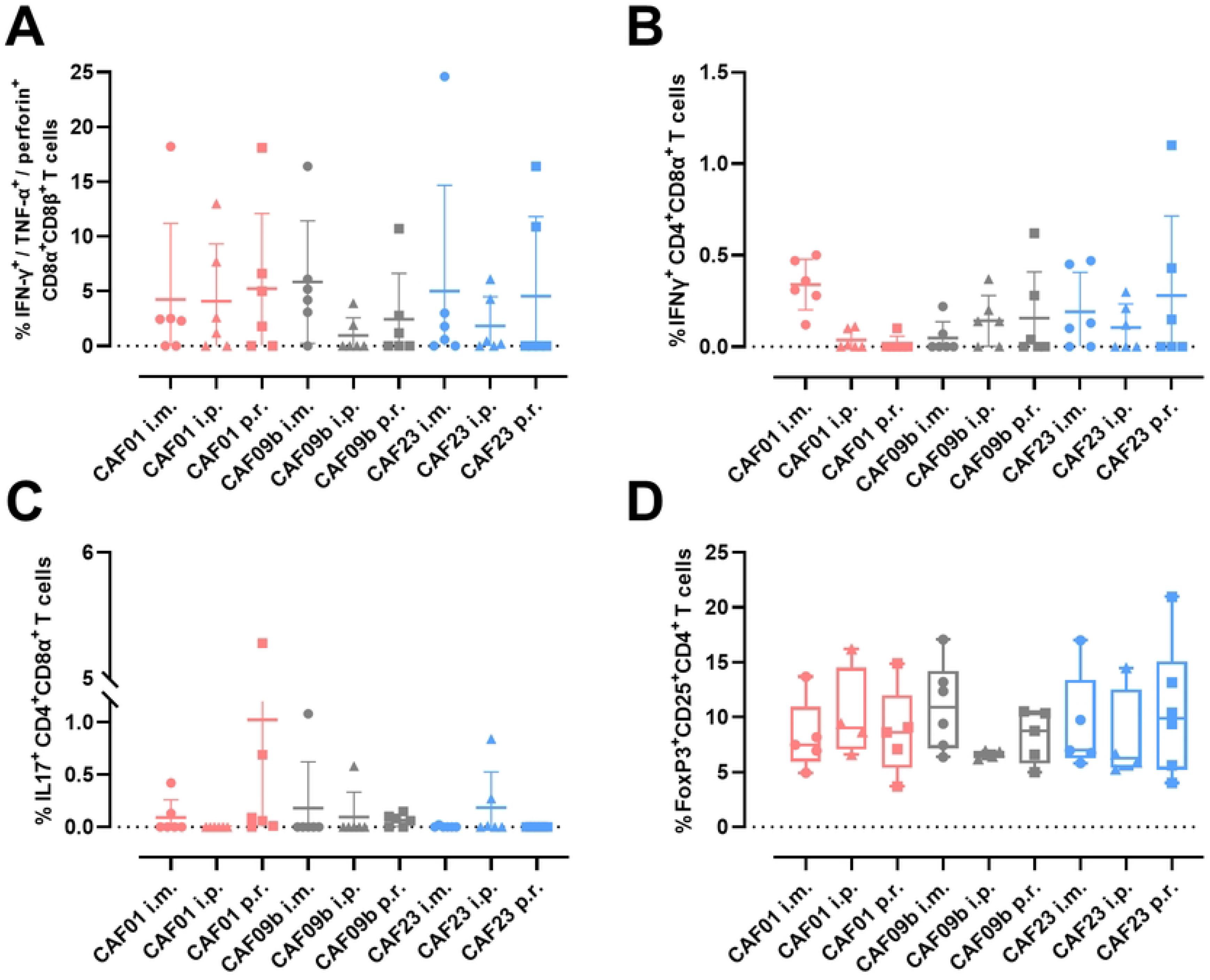
Cell-mediated immune responses grouped by cell type, measured by ICS flow cytometry: The background from NS controls was subtracted from the CTH522-stimulated samples, and the pigs with higher background than the signal after CTH22 re-stimulation had the Ag-specific response set to 0. Gating strategies and representative dot plots are shown in S1 and S2 Fig, respectively. A) Percentage of activated CTLs, defined by production of IFN-γ and/or TNF-α and/or perforin in the CD8α^+^CDβ^+^ T cell population across all groups after CTH522 re-stimulation. Percentage of CTH522-specific CD4^+^CD8α^+^ B) Th1 cells defined by IFN-γ production and C) Th17 cells defined by IL-17 production across all groups. D) Percentage of CD4^+^ Tregs expressing CD25 and activating FoxP3 transcription factor, shown in the NS controls. A) – D) Data represent two independent experiments, and all pigs (54) were initially included. No statistically significant differences were identified by two-way ANOVA. Data are presented as mean ± SD with data points representing responses from individual animals.

We also investigated gene expression of a few genes activated in CTLs. *TNFA* and *GZMB* expression did not significantly change with the adjuvant, though the Log_2_ RQ was around 4 in all groups (Fig 4A-B). However, CTH522 re-stimulation induced overexpression of *TNFA* in PBMCs of roughly half of the pigs in each adjuvant group (fold change > 0, on average 0.6 for CAF01). NKGD2 receptor complex encoded by *KLRK1* gene, was shown to be expressed on activated CD8^+^ T cells in mice, and human NK cells, CD8^+^ T cells and γδ T cells [34]. NKG2D can likely contribute to stimulation of proliferation and cytotoxicity in CD8^+^ T cells [35], which fits with our results where the *KLRK1* gene was upregulated by CTH522 re-stimulation in CAF09b group compared to CAF01 group (Fig 4A). This can be seen to a lower extent also on the RQ values (Fig 4B).

In the double-positive CD4^+^CD8α^+^ T cell population, IFN-γ production was overall the highest in the CAF01 i.m. group, (Fig 5B), which fits with our original hypothesis and previously published mouse studies [7]. Consistent IFN-γ production was also observed in the CAF09b i.p. and CAF23 i.m. groups, however, the variability within the groups was too high to see a clear pattern. In S2B Fig, representative flow cytometry dot plots of one animal from each group are displayed, showing IFN-γ^+^ CD4^+^CD8α^+^ population after CTH522 re-stimulation. Pigs possess high proportions of γδ T cells in the periphery, particularly young pigs where γδ T cells are the major T cell subset of PBMCs [32]. γδ T cells recognize unprocessed non-peptide Ags [36], therefore monitoring immune responses after protein Ag re-stimulation was not relevant. *IFNG* fold change expression between CTH522/NS (Fig 4C) and RQs (Fig 4D) between the adjuvant groups did not significantly differ. Nevertheless, *IFNG* was overexpressed (fold change > 0) across all the groups (Fig 4C). Similarly, *IL2* was overexpressed across all groups, but most profoundly in CAF01 group. IL-17 staining was very low, and majority of animals did not show any Ag-specific IL-17^+^ CD4^+^CD8α^+^ T cells (Fig 5C), but PHA stimulation induced low IL-17 responses, too (S4 Fig). The expression of *IL17A* and *RORC* encoding the transcription factor RORγt expressed by Th17 cells [37], did not indicate any differences between adjuvant groups, and all groups showed a Log_2_ fold change close to 0 for both genes, with some pigs having these genes downregulated (Fig 4E-F), supporting the results from flow cytometry staining. T regulatory (Treg) cells in pigs are in majority defined by CD4^+^ expression and are CD25^high^ expressing transcription factor FoxP3, but can express CD8α, too [38]. There was no difference in frequency of Treg cells between the groups (Fig 5D), which was confirmed by the lack of fold changes in *FOXP3, IL10* and *TGFB* gene expressions (Fig 4G) even though the RQ of *FOXP3* and *IL10* was between 3-4 (Fig 4H).

### 3.7. CAF23 increased expression of the central lymphoid homing marker CCR7

It has been shown previously that RA, a component of CAF23, can enhance the expression of integrin α4β7 for gut-specific homing in T cells upon activation [39]. There are no commercial pig-specific antibodies for homing markers and we therefore investigated gene expression of chosen homing markers in PBMCs. CCR7 orchestrates homing of T cells and Ag-presenting dendritic cells into the secondary lymphoid organs [40]. The *CCR7* gene was significantly upregulated between CTH522 re-stimulated and NS in the CAF23 group, compared to CAF01 group (p = 0.0037, Fig 6A) which fits with the rationale behind CAF23 design, but the RQs were at the same level for all groups (Fig 6B). On the other hand, no upregulating effect on *ITGA4* and *ITGB7* expression was detected in any of the groups (Fig 6A), even though the genes were being expressed (Fig 6B). CCR6 is a mucosal homing marker for Th17 cells and was shown to be induced by the CAF01 adjuvant in mice [5]. We could, however, not detect any changes in *CCR6* expression between the groups, whether taking fold changes or RQs into account (Fig 6A-B).

**Fig 6:**
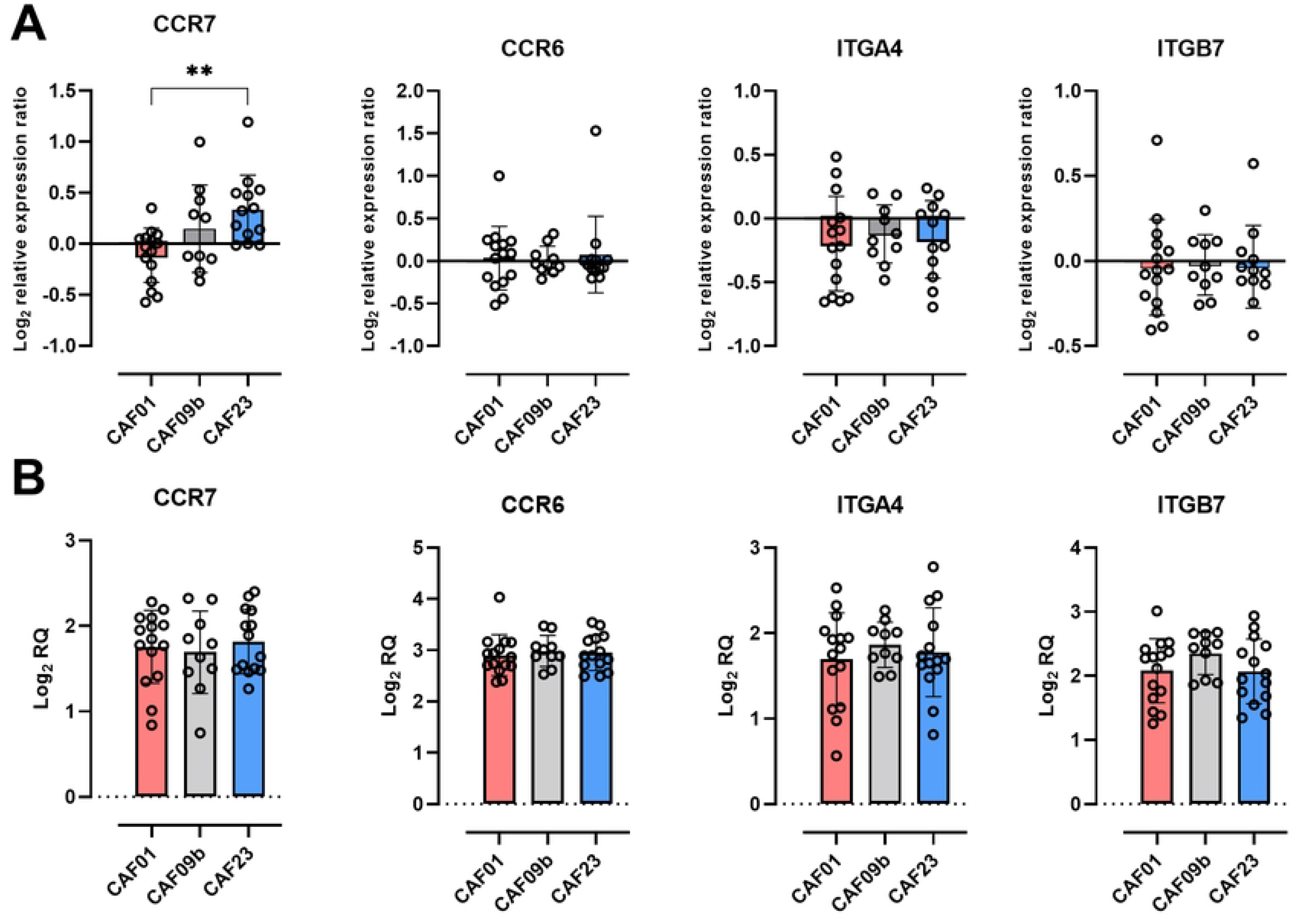
Expression of homing markers *CCR7*, *CCR6*, *ITGA4* and *ITGB7* in PBMCs measured by high-throughput qPCR: A) Log_2_ fold change expression ratio between CTH522 re-stimulated PBMCs and NS PBMCs and B) Log_2_ RQ. The different injection sites have been pooled under the respective adjuvant for the analysis. One outlier from the CAF09b i.p. group was excluded, due to low reproducibility between the cDNA replicates. Statistically significant differences are indicated by ** where p = 0.0037 (one-way ANOVA, followed by Tukey’s multiple comparisons test). There is no statistically significant difference, unless otherwise indicated. Data are presented as mean ± SD with data points representing responses from individual animals.

### 3.8. I.m. vaccine administration resulted in upregulation of multiple cytokine-encoding genes

Since we observed a significant effect of the injection site on anti-CTH522 antibody concentrations (Fig 2), we investigated, if this was translated into CMI-related gene expression too. The gene expression fold changes were therefore pooled into groups based on the injection site, disregarding the adjuvant. Most genes were not influenced by the site of injection with unchanged expression (fold change of 0), or the same order of upregulation across the groups. However, *IL17A* was significantly upregulated in the i.m. group compared to the p.r. group (p = 0.0430), where the results indicate downregulation (Fig 7). A similar, but non-significant trend was observed in *IFNG, IL2* and *TNFA* genes (Fig 7), which were upregulated to a higher extent in the i.m. group than in the i.p. and p.r. groups.

**Fig 7:**
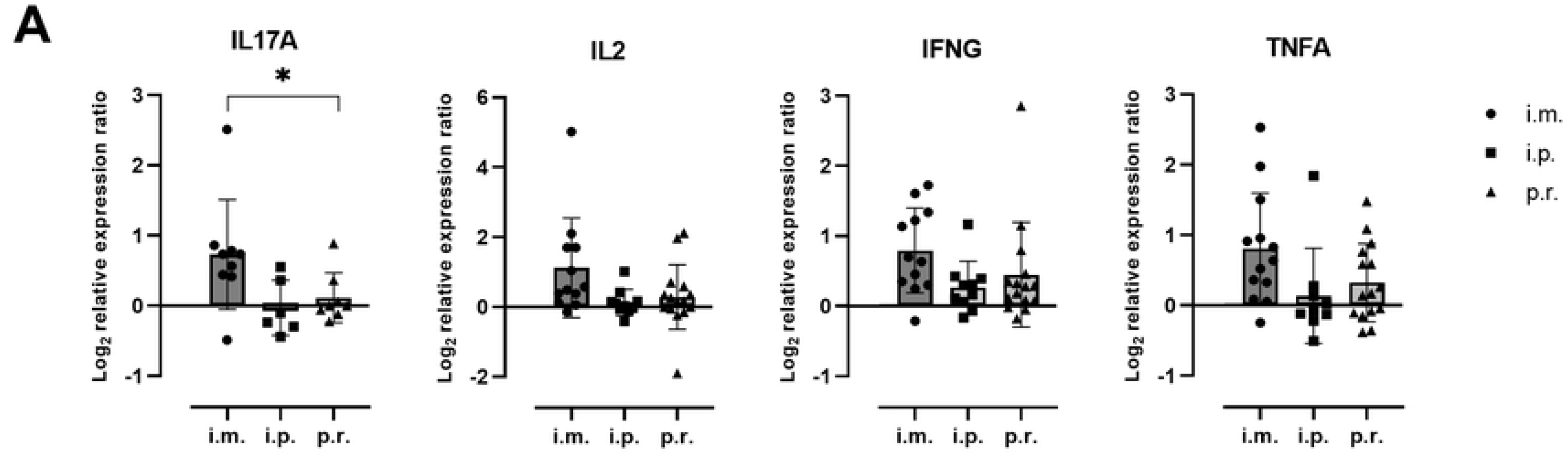
Expression of genes coding for cytokines grouped by injection sites measured by high-throughput qPCR: A) *IL17A*, *IL2*, *IFNG* and *TNFA* relative fold change expression between CTH522 re-stimulated PBMCs and NS PBMCs. The different adjuvants have been pooled under the respective injection sites for the analysis. One outlier from the CAF09b i.p. group was excluded, due to low reproducibility between the cDNA replicates. Statistically significant difference is indicated by *, where p = 0.0430 (one-way ANOVA, followed by Tukey’s multiple comparisons test). There is no statistically significant difference, unless otherwise indicated. Data are presented as mean ± SD with data points representing responses from individual animals.

### 3.9. IFN-γ release assays by ELISA and ELISpot confirm the flow cytometry results and variability

IFN-γ ELISA and ELISpot assays were performed in addition to ICS flow cytometry, but only 40 and 24 randomly chosen pigs, respectively, were included in these experiments. In ELISA, the concentration of IFN-γ secreted into the cell supernatant was measured, and the results followed the same trend as in ICS flow cytometry of IFN-γ in CTL or Th1 populations (Fig 5B and 8A). The amount of secreted IFN-γ ranged between pigs from non-responders to high-responders (concentration range after background subtraction was 1 – 1985 pg/mL), and yielded inconclusive results, as there were no differences between the groups and large variations within the groups. Similarly, as in the ICS, there was an *ex vivo* background IFN-γ interference in the NS controls (S5 Fig), therefore direct conclusions could not be made, as detecting Ag-specific responses was challenging.

**Fig 8:**
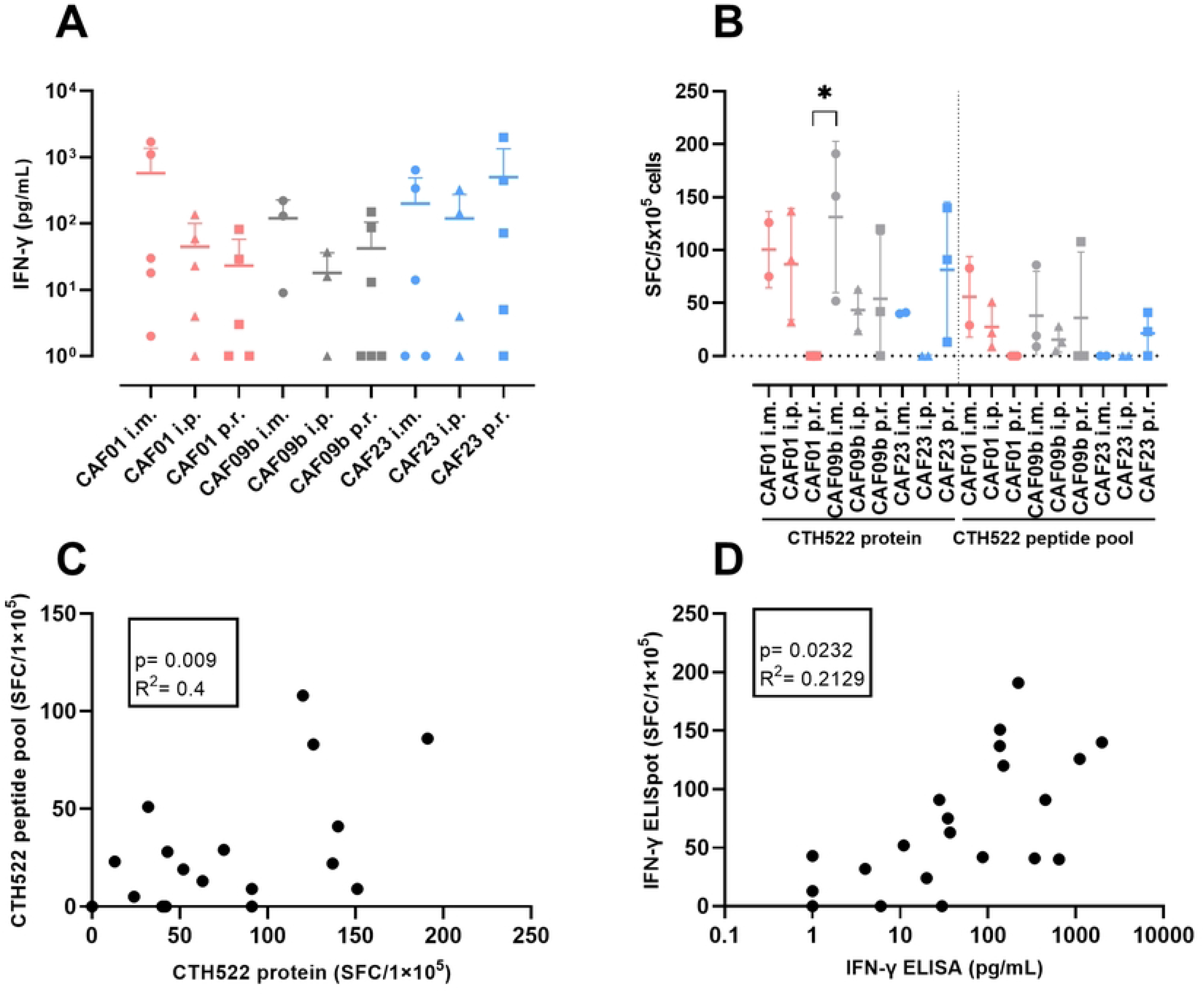
IFN-γ secretion quantified by ELISA and ELISpot: A) ELISA-based quantification of IFN-γ in cell supernatants after CTH522 re-stimulation. The background from NS controls was subtracted from the CTH522-stimulated, and all animals were assigned a minimal IFN-γ concentration of 1 pg/mL. B) Quantification of IFN-γ responses measured by ELISpot after re-stimulation by CTH522 protein (left) and predicted immunogenic 9- and 10-mer peptide pool (right). Data is presented as SFCs per 5×10^5^ PBMCs. The number of spots in NS controls was subtracted from the CTH522-stimulated samples and the pigs with negative amount of SFCs were set to 0 and considered non-responders. C) Correlation of SFCs count after CTH522 protein re-stimulation (x-axis) and CTH522 peptide pool re-stimulation (y-axis). D) Correlation of ELISA (x-axis) and ELISpot (y-axis) assays measuring IFN-γ secretion. Data in A) represent two independent experiments, and 40 randomly chosen pigs were initially included. No statistically significant differences were identified by two-way ANOVA. Data in B) and C) are from one experiment, and 24 randomly chosen pigs were initially included. Statistically significant difference is indicated by *, where p = 0.0348 (two-way ANOVA, followed by Tukey’s multiple comparisons test). There is no statistically significant difference, unless otherwise indicated. All data are presented as mean ± SD with data points representing responses from individual animals.

In the IFN-γ ELISpot assay, in addition to re-stimulation with the whole CTH522 protein, PBMCs were also stimulated with a pool of predicted 9- and 10-mer peptides from CTH522. This provided us a direct way of showing Ag-specific T cells (Fig 8B), as pigs in most groups were able to recognize these small peptides and produce IFN-γ to a higher level than the background (S5 Fig), while non-responders were assigned SFC count of 0. Even though the SFC counts were overall lower after peptide re-stimulation than after whole protein re-stimulation (on average 38 SFCs and 82 SFCs, respectively) it still verifies that these peptides were recognized. The CAF01 p.r. and CAF23 i.p. groups did not form higher number of SFCs than the background in neither protein, nor peptide pool stimulation, and the CAF23 i.m. group did not recognize the peptides. The CAF09b i.m. group produced the highest number of SFCs following whole CTH522 protein stimulation (Fig 8B). Even though the number of animals in each group is low, the IFN-γ production following whole protein versus peptide pool stimulation correlated significantly (R^2^ = 0.4; p = 0.0009; Fig 8C). The correlation between IFN-γ ELISA and ELISpot measurements was moderate (R^2^ = 0.2129; p = 0.0232; Fig 8D), which could be partially explained by a different nature of the assays. In the vast majority of pigs, the responders in ELISA correlated with the responders in ELISpot assay, however, in different magnitudes.

Even though both IFN-γ release assays provided inconclusive results, being in line with the ICS flow cytometry results, the SEB-stimulated positive controls produced on average 9-fold higher IFN-γ concentration in ELISA and 5-fold higher number of SFCs in ELISpot (S5 Fig), which supports the validity of the performed assays.

## 4. Discussion

This study offers new insights into immune responses to liposome-based adjuvants in a large animal model, the pig, which contributes to the well-characterized knowledge on CAF adjuvants we have from mouse model. Here, we showed the induction of antibody and CMI responses when administering CTH522 Ag formulated with either CAF01, CAF09b, CAF16 or CAF23 adjuvants via the i.m., i.p. or p.r. route in commercial pigs housed under field conditions. We showed that immunizations with these adjuvants consistently induce high titers of IgA and IgG antibodies and variable production of IFN-γ after CTH522 vaccine Ag re-stimulation. As now well-acknowledged, inbred laboratory mice living in sterile environment bear a less representative immune system, which for example lacks differentiated memory CD8^+^ T cell subsets, resembling more neonates than adult human population in this respect [41], [42]. The aim of this study was to test different administration regimes and vaccine adjuvants side-by-side in pigs, and to obtain additional knowledge on effects of adjuvant immunomodulators and injection site in a large animal model under field conditions.

A common interest in fine-tuning the immune responses lead to development of adjuvants, such as the CAF platform, polarizing the immune response towards the desired type of immunity [5], [43]. All adjuvants induced significant humoral and CMI responses, however, a polarizing effect, or differences between the individual adjuvants were lacking in pigs. Despite all the precautions that have been taken, such as choosing seemingly healthy animals and immediate handling of the blood samples, unspecific *ex vivo* cytokine background was very high and detected by the sensitive ICS flow cytometry and IFN-γ release assays in most pigs (S3 – 5 Fig). One of the possible explanations is that pigs received a standard circovirus immunization just a week before the start of our experiment. Due to this and possibly other reasons, detecting the rare Ag-specific T cells became challenging. As we could not identify clear-cut CMI responses in the initial experiment, we decided to add another dimension to our study and investigate gene expression in PBMCs by microfluidic high-throughput qPCR. However, using this method, we were not able to find different immune profiles of the tested adjuvants, either.

Overall, CAF01 administered i.m. yielded the most consistent results in terms of antibody secretion and CMI, particularly IFN-γ. This evidence is further supported by the gene expression analysis, where CAF01 animals re-stimulated with CTH522 were clustering separately from the rest of the CTH522 and NS groups, suggesting a differential gene expression, clustering closer to the positive SEB control. CAF01 formulated with different *C. trachomatis* Ags (Hirep1 and CTH93), administered as i.m. prime and intranasal boost, has previously been tested in Göttingen minipigs, where significant systemic IgG, mucosal SIgA and cell-mediated IFN-γ and IL-17 responses were induced [10]. This is partially in line with our results, but the minipigs in the previous study were free of specific pathogens, older and housed in clean isolation units. Additionally, mucosal IgA responses were triggered both with or without the adjuvant, pointing to the mucosal administration effect of intranasal boost immunization, which was not applied in the current study. It is of note that IgA after subcutaneous CAF01 immunization is practically absent in mice studies, and arises both systematically and locally only after subcutaneous prime followed by intranasal boost [31] or when RA is incorporated into the adjuvant, as observed in CB6F1 (C57BL/6xBALB/c) mice [17]. In pigs, serum IgA was present after parental immunization with all CAF adjuvants, irrespective of the injection site. The apparent difference in serum IgA response to standard adjuvants between pigs and mice may have obliterated the RA adjuvant effect to induce IgA and SIgA response in serum and mucus of pigs, as CAF16 or CAF23 did not differ from CAF01 and CAF09b in this ability.

CAF09 administered i.p. has previously been shown to induce CTLs in Göttingen minipigs after repeated immunizations, but CAF09 was not compared to any other adjuvant [23]. Although we did not see any difference in CTL response with varying adjuvants, the magnitude of T cell responses measured by ELISpot and ICS in the current study, are comparable to the responses acquired in the previous study. In our study, i.p. administration of CAF09b was not significantly better at inducing CTLs, than i.m. or p.r. injections. The i.p. immunizations were not performed ultrasound-guided [15], so the injection could have been mistakenly administered into the intestinal lumen, explaining the lack of response in some pigs.

Pigs are susceptible to several species from the *Chlamydiaceae* family, most common being *C. suis,* which was found in 95.7% of fecal swabs in Swiss fattening pigs [44]. There are indications of cross-protection between *C. trachomatis* and *C. suis*, as CMI responses were detected after both, homologous or heterologous re-stimulation [45]. In our study, we have not tested the animals for chlamydial DNA content, but since there were no anti-CTH522 antibodies in serum at day 0 (Figs 1 and 2), the pigs were considered chlamydia-naïve. However, we cannot completely rule out that the piglets or sows have previously encountered *Chlamydiaceae* strains. If so, the vaccines could be regarded as post-exposure immunizations, which could influence the resulting immune responses.

The impact of vaccine-site microenvironment is not a new concept and a recent study found that the injection site is also an important site of Ag presentation, where T cell responses can be generated locally, independent of the draining lymph node after subcutaneous/intradermal immunizations [46]. Ischiorectal fossa is commonly used as injection site in cattle for prostaglandin, but has not been evaluated as an injection site for prophylactic vaccines [20]. Here, we hypothesized that the p.r. injection would drain to mucosa-associated *Anorectales* lymph nodes and facilitate an enhanced effect of RA incorporation resulting in higher IgA titers. In contrast, the p.r. injection route was shown to be the least effective in regards to potentiating serum IgA and IgG responses, while in mucus all injection sites induced roughly same levels of IgA and SIgA. If injection into p.r. space is not performed correctly, it can be wasted into the rectum [20], which is likely an explanation for several low-responding pigs in the p.r. groups, but does not explain the overall lower response to vaccination in this group. This observation might reflect a lower number of circulating vaccine-activated T cells after p.r. immunization compared to i.m. or i.p. immunizations. The i.m. injection site, irrespective on the adjuvant, was more prone to stimulate cytokines’ gene expression and therefore overall could have been more immunogenic.

In this study we found that injection sites, but not immunomodulation of liposome-based adjuvants, had a significant impact on Ag-specific antibodies in serum, while neither injection site, nor adjuvant significantly affected the CMI responses, which could partly be due to variable and high cytokine background in these farm animals. These results indicate that when using pigs in the real-life setting, immunomodulatory adjuvants that induce well-defined immune responses in e.g. mouse models, may not perform similarly in a standard prime-boost immunization scheme. Traditionally, animal vaccines have employed emulsion-based oil-in-water/water-in-oil adjuvants [47] and there is evidence suggesting that different types of emulsions represent a preferred adjuvant in veterinary vaccines, because they induce strong immune responses, both short- and long-term, and are usually well-tolerated [48]. The use of liposome adjuvants in pigs is scarcely published, and may not be optimal adjuvant formulation for pigs. Furthermore, it can be difficult to predict the best animal model for translating into humans, and our study shows the limitations of using field-setting pigs for human adjuvant development. While non-human primates may offer better predictability of adjuvant effects in humans, direct comparison of different adjuvant systems in clinical trials is still necessary. Overall, our study highlights that responses to vaccination with different adjuvants vary dramatically between animal species, indicating that for veterinary vaccines, adjuvant comparison studies need to be performed in their target species early on in development.

## Acknowledgments

We would like to acknowledge Karin Tarp for help with microfluidic high-throughput qPCR. Animal caretakers are thanked for taking good care of the pigs.

## Author Contributions

The authors confirm contribution to the paper as follows: study conception and design: EŠ, GJ, GE; data collection: EŠ, JTJ; analysis and interpretation of results: EŠ, KS, GJ, GE; draft manuscript preparation: EŠ, GJ, GE, GKP. All authors read and approved the final version of the manuscript.

## Supporting information

**S1 Table:**
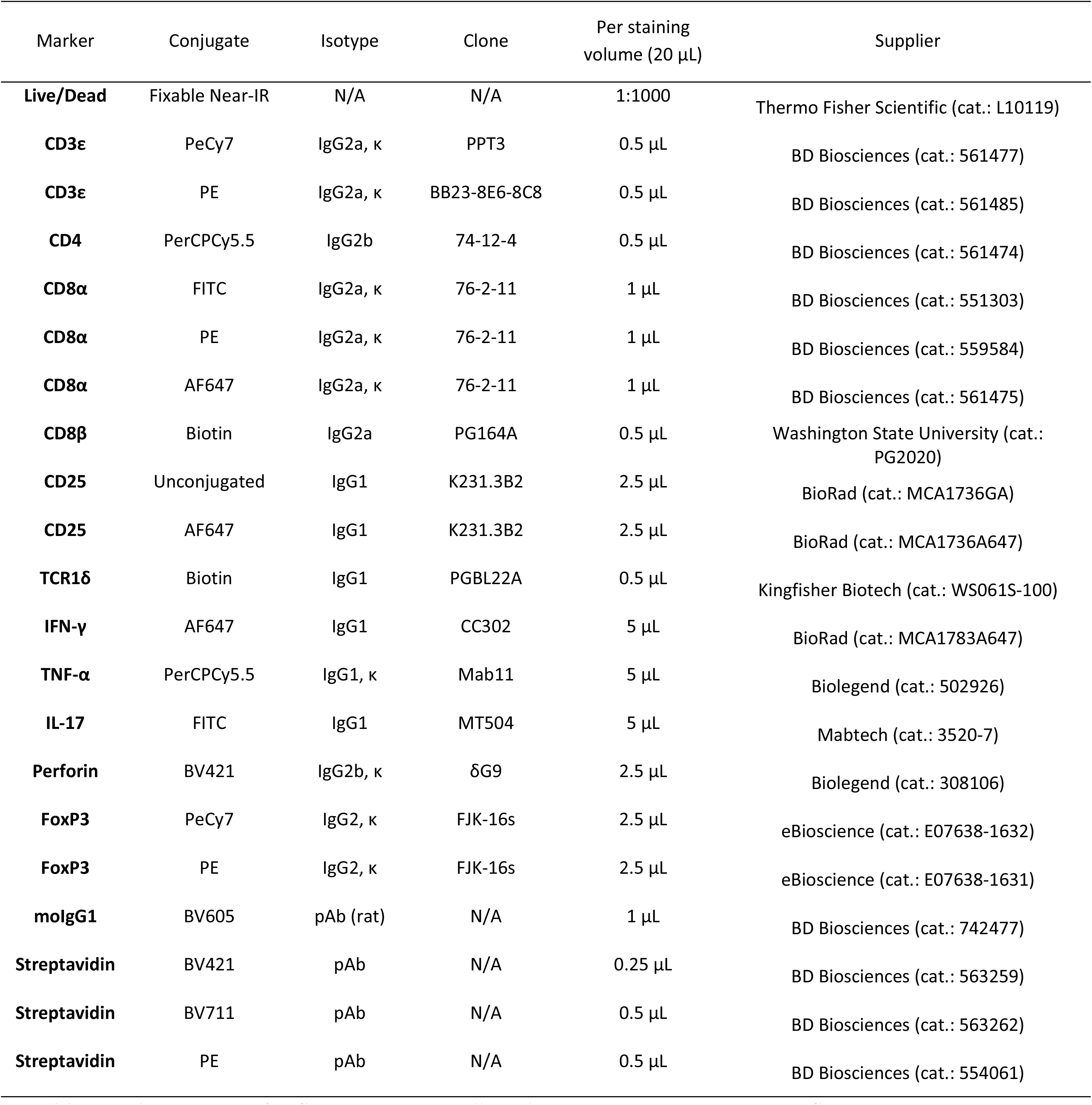
Antibodies used for flow cytometry. All antibodies were titrated prior to flow cytometry staining. All antibodies are monoclonal (mAb), with mouse as the host species, unless otherwise stated (polyclonal – pAb).

**S2 Table:**
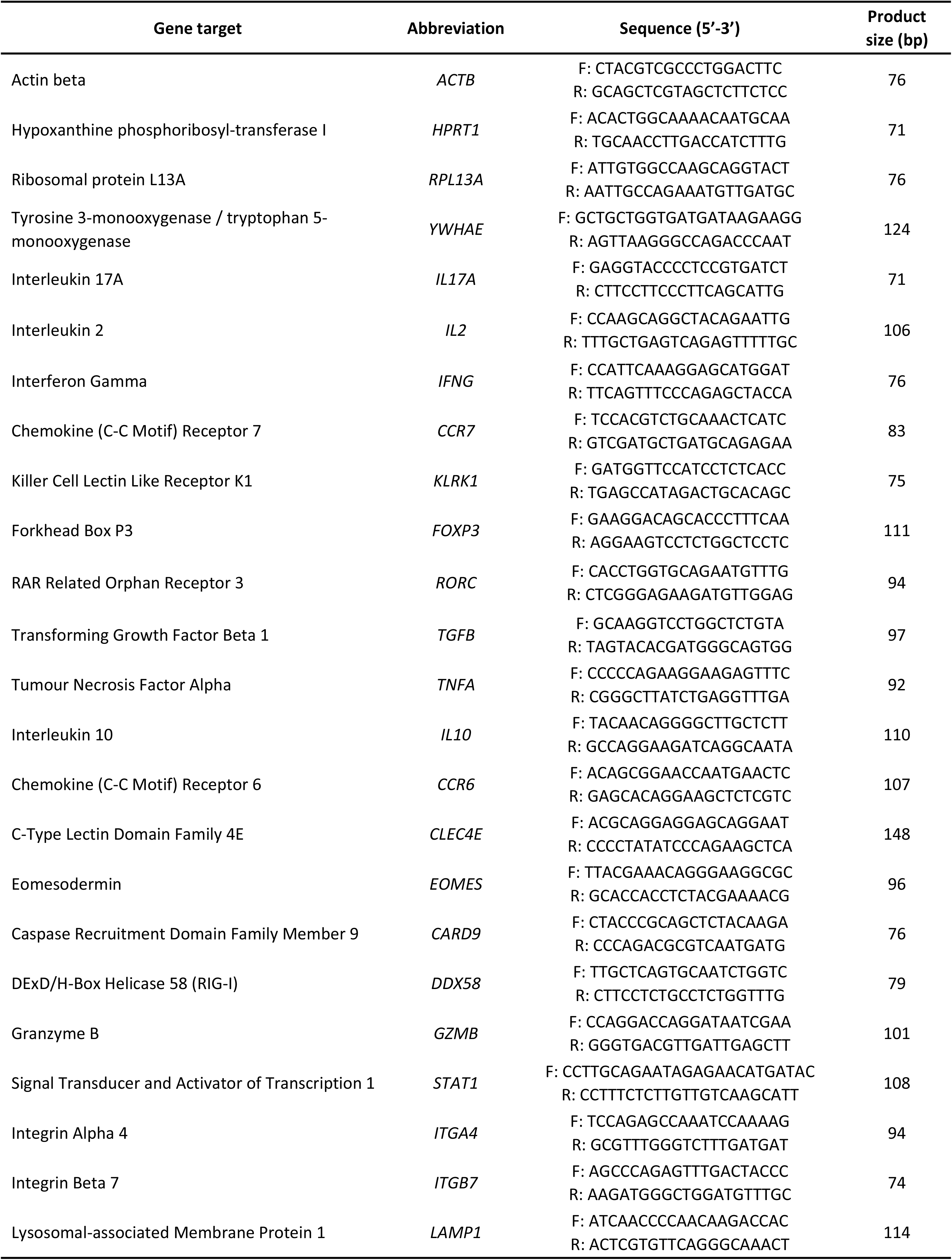
List of primers used for high throughput RT-qPCR.

**S1 Fig: Representative gating strategy used for flow cytometry.** For analysis of flow cytometric data, cells were first gated on viable cells negative for LIVE/DEAD™ Fixable Near-IR amine reactive dye; single cells were then selected based on the FSC-A/FSC-H, followed by lymphocytes population based on FSC-A/SSC-A. T cells were gated based on CD3^+^ staining. The final analysis for each panel was as following: A) CTL panel: CD3^+^ T cells were gated on CD8α^+^ T cells and subsequently gated on CD8β^+^ and CD8β^-^. Cytokine positive cells were detected as IFN-γ^+^CD8β^+^, TNF-α^+^CD8β^+^ and perforin^+^CD8β^+^ T cells. B) Th1/Th17 panel: CD3^+^ T cells were gated on TCR1δ marker, TCR1δ^-^T cells were further gated based on CD4/CD8α markers and double-positive CD4^+^CD8α^+^ cells were gated on IFN-γ+ and IL-17^+^. C) Treg panel: CD3^+^ T cells were gated on CD4/CD8α markers, and CD4^+^ T cells were further gated on CD25^+^ and FoxP3^+^.

**S2 Fig: IFN-γ production in response to CTH522 Ag.** A) Representative flow cytometry dot plots of IFN-γ^+^ CD3^+^ Th1 cells across all groups. Numbers indicate the percentage of IFN-γ^+^ cells as a proportion of double positive CD4^+^CD8α^+^ T cells, pre-gated on live cells – single cells – lymphocytes and CD3^+^ T cells. B) Representative flow cytometry dot plots of IFN-γ^+^ CD3^+^ CTL cells across all groups. Numbers indicate the percentage of IFN-γ^+^ cells as a proportion of double positive CD8α^+^CD8β^+^ T cells, pre-gated on live cells – single cells – lymphocytes and CD3^+^ T cells.

**S3 Fig: Positive and negative controls in the IFN-γ and IL-17 ELISA assays and ICS flow cytometry (initial experiment).** A) SEB-stimulated positive control and NS negative control of IFN-γ and B) IL-17 cytokine release assays. All animals were assigned a minimal IFN-γ and IL-17 concentration of 1 pg/mL. Percentage of C) IFN-γ-producing Th1 cells and D) IL-17-producing Th17 cells after PHA stimulation and NS in media shown as proportion of CD4^+^CD8α^+^ T cells. Data are presented as mean ± SD with data points representing responses from individual animals

**S4 Fig: Positive and negative controls in ICS flow cytometry (main experiment):** A) Percentage of activated CTLs, defined by production of IFN-γ and/or TNF-α and/or perforin in the CD8α^+^CDβ^+^ T cell population across all groups after PHA stimulation (left), PHA stimulation after background signal subtraction (middle) and in media (right). Percentage of B) IFN-γ-producing Th1 cells and C) IL-17-producing Th17 cells after PHA stimulation (left), PHA stimulation after background signal subtraction (middle) and in media (right) shown as proportion of CD4^+^CD8α^+^ T cells. Data are presented as mean ± SD with data points representing responses from individual animals.

**S5 Fig: Positive and negative controls in the IFN-γ ELISA and IFN-γ ELISpot assays.** A) SEB-stimulated positive control (left) and NS negative control (right) of IFN-γ release assay. All animals were assigned a minimal IFN-γ and IL-17 concentration of 1 pg/mL. B) SFCs per 5×10^5^ cells in SEB-stimulated positive control (left) and NS negative control (right). C) IFN-γ ELISpot images in triplicates from one representative animal in response to stimulation with SEB, media (NS), CTH522 protein and CTH522 peptides. Data are presented as mean ± SD with data points representing responses from individual animals.

**S6 Fig: Expression of genes characteristic for T cell activation, pathways activated by CAF01 and CAF09b, respectively measured by high-throughput qPCR:** A) *EOMES* and *LAMP1* Log_2_ fold change, and B) Log_2_ RQ in the CTH522-re-stimulated PBMCs. C) *CLEC4E* and *CARD9* Log_2_ fold change, and D) Log_2_ RQ in the CTH522-re-stimulated PBMCs. E) *DDX58* and *STAT1* A) Log_2_ fold change, and F) Log_2_ RQ in the CTH522 re-stimulated PBMCs. Fold changes are represented as a ratio between relative gene quantity in CTH522 re-stimulated PBMCs and NS PBMCs. The different injection sites were pooled under the respective adjuvant. Data are presented as mean ± SD with data points representing responses from individual animals.

## References

[1] S. Nooraei, A. Sarkar Lotfabadi, M. Akbarzadehmoallemkolaei, and N. Rezaei, “Immunogenicity of Different Types of Adjuvants and Nano-Adjuvants in Veterinary Vaccines: A Comprehensive Review,” Vaccines (Basel), vol. 11, no. 2, p. 453, Feb. 2023, doi: 10.3390/vaccines11020453.

[2] S. G. Reed, M. T. Orr, and C. B. Fox, “Key roles of adjuvants in modern vaccines,” Nat Med, vol. 19, no. 12, pp. 1597–1608, Dec. 2013, doi: 10.1038/nm.3409.

[3] R. L. Coffman, A. Sher, and R. A. Seder, “Vaccine adjuvants: putting innate immunity to work,” Immunity, vol. 33, no. 4, pp. 492–503, Oct. 2010, doi: 10.1016/j.immuni.2010.10.002.

[4] B. Pulendran, P. S Arunachalam, and D. T. O’Hagan, “Emerging concepts in the science of vaccine adjuvants,” Nat Rev Drug Discov, vol. 20, no. 6, pp. 454–475, Jun. 2021, doi: 10.1038/s41573-021-00163-y.

[5] G. K. Pedersen, P. Andersen, and D. Christensen, “Immunocorrelates of CAF family adjuvants,” Semin Immunol, vol. 39, pp. 4–13, Oct. 2018, doi: 10.1016/j.smim.2018.10.003.

[6] G. K. Pedersen, K. Wørzner, P. Andersen, and D. Christensen, “Vaccine Adjuvants Differentially Affect Kinetics of Antibody and Germinal Center Responses,” Front Immunol, vol. 11, p. 579761, 2020, doi: 10.3389/fimmu.2020.579761.

[7] J. Davidsen et al., “Characterization of cationic liposomes based on dimethyldioctadecylammonium and synthetic cord factor from M. tuberculosis (trehalose 6,6’-dibehenate)-a novel adjuvant inducing both strong CMI and antibody responses,” Biochim Biophys Acta, vol. 1718, no. 1–2, pp. 22–31, Dec. 2005, doi: 10.1016/j.bbamem.2005.10.011.

[8] H. Schoenen et al., “Cutting Edge: Mincle Is Essential for Recognition and Adjuvanticity of the Mycobacterial Cord Factor and its Synthetic Analog Trehalose-Dibehenate,” The Journal of Immunology, vol. 184, no. 6, pp. 2756–2760, Mar. 2010, doi: 10.4049/jimmunol.0904013.

[9] K. Werninghaus et al., “Adjuvanticity of a synthetic cord factor analogue for subunit Mycobacterium tuberculosis vaccination requires FcRγ–Syk–Card9–dependent innate immune activation,” Journal of Experimental Medicine, vol. 206, no. 1, pp. 89–97, Jan. 2009, doi: 10.1084/jem.20081445.

[10] E. Lorenzen et al., “Intramuscular Priming and Intranasal Boosting Induce Strong Genital Immunity Through Secretory IgA in Minipigs Infected with Chlamydia trachomatis,” Front Immunol, vol. 6, p. 628, 2015, doi: 10.3389/fimmu.2015.00628.

[11] E. Lorenzen et al., “Multi-component prime-boost Chlamydia trachomatis vaccination regimes induce antibody and T cell responses and accelerate clearance of infection in a non-human primate model,” Front Immunol, vol. 13, p. 1057375, 2022, doi: 10.3389/fimmu.2022.1057375.

[12] S. Abraham et al., “Safety and immunogenicity of the chlamydia vaccine candidate CTH522 adjuvanted with CAF01 liposomes or aluminium hydroxide: a first-in-human, randomised, double-blind, placebo-controlled, phase 1 trial,” The Lancet Infectious Diseases, vol. 19, no. 10, pp. 1091–1100, Oct. 2019, doi: 10.1016/S1473-3099(19)30279-8.

[13] R. Billeskov et al., “Testing the H56 Vaccine Delivered in 4 Different Adjuvants as a BCG-Booster in a Non-Human Primate Model of Tuberculosis,” PLoS One, vol. 11, no. 8, p. e0161217, 2016, doi: 10.1371/journal.pone.0161217.

[14] K. S. Korsholm et al., “Induction of CD8+ T-cell responses against subunit antigens by the novel cationic liposomal CAF09 adjuvant,” Vaccine, vol. 32, no. 31, pp. 3927–3935, Jun. 2014, doi: 10.1016/j.vaccine.2014.05.050.

[15] S. K. Mørk et al., “Personalized therapy with peptide-based neoantigen vaccine (EVX-01) including a novel adjuvant, CAF®09b, in patients with metastatic melanoma,” Oncoimmunology, vol. 11, no. 1, p. 2023255, 2022, doi: 10.1080/2162402X.2021.2023255.

[16] G. V. Long et al., “KEYNOTE – D36: personalized immunotherapy with a neoepitope vaccine, EVX-01 and pembrolizumab in advanced melanoma,” Future Oncology, vol. 18, no. 31, pp. 3473–3480, Oct. 2022, doi: 10.2217/fon-2022-0694.

[17] D. Christensen et al., “A Liposome-Based Adjuvant Containing Two Delivery Systems with the Ability to Induce Mucosal Immunoglobulin A Following a Parenteral Immunization,” ACS Nano, vol. 13, no. 2, pp. 1116–1126, 26 2019, doi: 10.1021/acsnano.8b05209.

[18] S. T. Schmidt et al., “The administration route is decisive for the ability of the vaccine adjuvant CAF09 to induce antigen-specific CD8+ T-cell responses: The immunological consequences of the biodistribution profile,” Journal of Controlled Release, vol. 239, pp. 107–117, Oct. 2016, doi: 10.1016/j.jconrel.2016.08.034.

[19] H. Jin et al., “Higher immune response induced by vaccination in Houhai acupoint relates to the lymphatic drainage of the injection site,” Res Vet Sci, vol. 130, pp. 230–236, Jun. 2020, doi: 10.1016/j.rvsc.2020.03.018.

[20] S. C. Holland, W. D. Whittier, S. G. Clark, S. A. Hafez, and W. S. Swecker, “Comparison of luteolysis and timed artificial insemination pregnancy rates after administration of PGF2α in the muscle or the ischiorectal fossa in cattle,” Animal Reproduction Science, vol. 198, pp. 11–19, Nov. 2018, doi: 10.1016/j.anireprosci.2018.07.003.

[21] V. Gerdts et al., “Large animal models for vaccine development and testing,” ILAR J, vol. 56, no. 1, pp. 53–62, 2015, doi: 10.1093/ilar/ilv009.

[22] T. Käser, “Swine as biomedical animal model for T-cell research-Success and potential for transmittable and non-transmittable human diseases,” Mol Immunol, vol. 135, pp. 95–115, Jul. 2021, doi: 10.1016/j.molimm.2021.04.004.

[23] N. H. Overgaard, T. M. Frøsig, J. T. Jakobsen, S. Buus, M. H. Andersen, and G. Jungersen, “Low antigen dose formulated in CAF09 adjuvant Favours a cytotoxic T-cell response following intraperitoneal immunization in Göttingen minipigs,” Vaccine, vol. 35, no. 42, pp. 5629–5636, 09 2017, doi: 10.1016/j.vaccine.2017.08.057.

[24] L. E. Pedersen, G. Jungersen, M. R. Sorensen, C.-S. Ho, and D. F. Vadekær, “Swine Leukocyte Antigen (SLA) class I allele typing of Danish swine herds and identification of commonly occurring haplotypes using sequence specific low and high resolution primers,” Vet. Immunol. Immunopathol., vol. 162, no. 3–4, pp. 108–116, Dec. 2014, doi: 10.1016/j.vetimm.2014.10.007.

[25] S. Meling, K. Skovgaard, K. Bårdsen, P. M. Helweg Heegaard, and M. J. Ulvund, “Expression of selected genes isolated from whole blood, liver and obex in lambs with experimental classical scrapie and healthy controls, showing a systemic innate immune response at the clinical end-stage,” BMC Vet Res, vol. 14, no. 1, p. 281, Sep. 2018, doi: 10.1186/s12917-018-1607-9.

[26] E. M. Agger et al., “Cationic Liposomes Formulated with Synthetic Mycobacterial Cordfactor (CAF01): A Versatile Adjuvant for Vaccines with Different Immunological Requirements,” PLoS One, vol. 3, no. 9, p. e3116, Sep. 2008, doi: 10.1371/journal.pone.0003116.

[27] J. S. Woodworth, et al., “A novel adjuvant formulation induces robust Th1/Th17 memory and mucosal recall responses in Non-Human Primates,” bioRxiv, p. 2023.02.23.529651, Feb. 2023, doi: 10.1101/2023.02.23.529651.

[28] W. Gerner, S. C. Talker, H. C. Koinig, C. Sedlak, K. H. Mair, and A. Saalmüller, “Phenotypic and functional differentiation of porcine αβ T cells: Current knowledge and available tools,” Molecular Immunology, vol. 66, no. 1, pp. 3–13, Jul. 2015, doi: 10.1016/j.molimm.2014.10.025.

[29] N. Wang et al., “Primary characterization of the immune response in pigs infected with Trichinella spiralis,” Vet Res, vol. 51, p. 17, 2020, doi: 10.1186/s13567-020-0741-0.

[30] Y. Li, L. Jin, and T. Chen, “The Effects of Secretory IgA in the Mucosal Immune System,” Biomed Res Int, vol. 2020, p. 2032057, 2020, doi: 10.1155/2020/2032057.

[31] D. Christensen, R. Mortensen, I. Rosenkrands, J. Dietrich, and P. Andersen, “Vaccine-induced Th17 cells are established as resident memory cells in the lung and promote local IgA responses,” Mucosal Immunol, vol. 10, no. 1, pp. 260–270, Jan. 2017, doi: 10.1038/mi.2016.28.

[32] H. Yang and R. M. E. Parkhouse, “Phenotypic classification of porcine lymphocyte subpopulations in blood and lymphoid tissues,” Immunology, vol. 89, no. 1, pp. 76–83, 1996, doi: 10.1046/j.1365-2567.1996.d01-705.x.

[33] H. Yang and R. M. Parkhouse, “Differential expression of CD8 epitopes amongst porcine CD8-positive functional lymphocyte subsets,” Immunology, vol. 92, no. 1, pp. 45–52, Sep. 1997, doi: 10.1046/j.1365-2567.1997.00308.x.

[34] S. Bauer et al., “Activation of NK Cells and T Cells by NKG2D, a Receptor for Stress-Inducible MICA,” Science, vol. 285, no. 5428, pp. 727–729, Jul. 1999, doi: 10.1126/science.285.5428.727.

[35] L. I. R. Ehrlich et al., “Engagement of NKG2D by cognate ligand or antibody alone is insufficient to mediate costimulation of human and mouse CD8+ T cells,” J Immunol, vol. 174, no. 4, pp. 1922–1931, Feb. 2005, doi: 10.4049/jimmunol.174.4.1922.

[36] Y. Tanaka et al., “Nonpeptide ligands for human gamma delta T cells,” Proc Natl Acad Sci U S A, vol. 91, no. 17, pp. 8175–8179, Aug. 1994, doi: 10.1073/pnas.91.17.8175.

[37] I. I. Ivanov et al., “The orphan nuclear receptor RORgammat directs the differentiation program of proinflammatory IL-17+ T helper cells,” Cell, vol. 126, no. 6, pp. 1121–1133, Sep. 2006, doi: 10.1016/j.cell.2006.07.035.

[38] T. Käser, W. Gerner, S. E. Hammer, M. Patzl, and A. Saalmüller, “Detection of Foxp3 protein expression in porcine T lymphocytes,” Veterinary Immunology and Immunopathology, vol. 125, no. 1, pp. 92–101, Sep. 2008, doi: 10.1016/j.vetimm.2008.05.007.

[39] M. Iwata, A. Hirakiyama, Y. Eshima, H. Kagechika, C. Kato, and S.-Y. Song, “Retinoic acid imprints gut-homing specificity on T cells,” Immunity, vol. 21, no. 4, pp. 527–538, Oct. 2004, doi: 10.1016/j.immuni.2004.08.011.

[40] R. Förster, A. C. Davalos-Misslitz, and A. Rot, “CCR7 and its ligands: balancing immunity and tolerance,” Nat Rev Immunol, vol. 8, no. 5, Art. no. 5, May 2008, doi: 10.1038/nri2297.

[41] L. K. Beura et al., “Normalizing the environment recapitulates adult human immune traits in laboratory mice,” Nature, vol. 532, no. 7600, pp. 512–516, Apr. 2016, doi: 10.1038/nature17655.

[42] L. Cai, H. Xu, and Z. Cui, “Factors Limiting the Translatability of Rodent Model-Based Intranasal Vaccine Research to Humans,” AAPS PharmSciTech, vol. 23, no. 6, p. 191, Jul. 2022, doi: 10.1208/s12249-022-02330-9.

[43] A. R. Spickler and J. A. Roth, “Adjuvants in veterinary vaccines: modes of action and adverse effects,” J Vet Intern Med, vol. 17, no. 3, pp. 273–281, 2003, doi: 10.1111/j.1939-1676.2003.tb02448.x.

[44] K. Hoffmann et al., “Prevalence of Chlamydial Infections in Fattening Pigs and Their Influencing Factors,” PLOS ONE, vol. 10, no. 11, p. e0143576, Nov. 2015, doi: 10.1371/journal.pone.0143576.

[45] T. Käser et al., “Chlamydia suis and Chlamydia trachomatis induce multifunctional CD4 T cells in pigs,” Vaccine, vol. 35, no. 1, pp. 91–100, Jan. 2017, doi: 10.1016/j.vaccine.2016.11.050.

[46] M. O. Meneveau, P. Kumar, K. T. Lynch, S. P. Patel, and C. L. Slingluff, “The vaccine-site microenvironment: impacts of antigen, adjuvant, and same-site vaccination on antigen presentation and immune signaling,” J Immunother Cancer, vol. 10, no. 3, p. e003533, Mar. 2022, doi: 10.1136/jitc-2021-003533.

[47] Y. Burakova, R. Madera, S. McVey, J. R. Schlup, and J. Shi, “Adjuvants for Animal Vaccines,” Viral Immunology, vol. 31, no. 1, pp. 11–22, Jan. 2018, doi: 10.1089/vim.2017.0049.

[48] J. Aucouturier, L. Dupuis, and V. Ganne, “Adjuvants designed for veterinary and human vaccines,” Vaccine, vol. 19, no. 17–19, pp. 2666–2672, Mar. 2001, doi: 10.1016/S0264-410X(00)00498-9.

